# Developmental Reactive Oxygen Species Target Histone Methylation to Individualize Stress resistance and Lifespan

**DOI:** 10.1101/756742

**Authors:** Daphne Bazopoulou, Daniela Knoefler, Yongxin Zheng, Kathrin Ulrich, Bryndon Oleson, Lihan Xie, Minwook Kim, Anke Kaufmann, Young-Tae Lee, Yali Dou, Yong Chen, Shu Quan, Ursula Jakob

## Abstract

A central aspect of aging research concerns the question as to when individuality in lifespan arises and what mechanism(s) promote and potentially manifest individual differences in longevity. We have now discovered that a transient increase in reactive oxygen species (ROS), which occurs naturally during early development in a subpopulation of synchronized *Caenorhabditis elegans*, sets processes into motion that increase stress resistance, improve redox homeostasis and ultimately prolong lifespan in those animals. We find that these effects are linked to the global ROS-mediated decrease in developmental histone H3K4me3 levels. Studies in HeLa cells confirmed that global H3K4me3 levels are ROS-sensitive, and that depletion of H3K4me3 levels increases stress resistance in mammalian cell cultures. *In vitro* studies identified the Set1-MML histone methyltransferase as the redox sensitive unit of the H3K4-trimethylating COMPASS complex. Our findings imply a novel link between early-life events, ROS-sensitive epigenetic marks, stress resistance and lifespan.

---

It is well established that genetic effects account for less than 25% of the observed differences in human lifespan ^1,2^. However, the remaining differences are not entirely attributable to environmental factors either. For instance, even when isogenic animals, such as *Caenorhabditis elegans*, are cultivated under identical environmental conditions, individual lifespans can vary by over 50-fold ^3–5^. These results suggest that other, more stochastic factors account for lifespan variations and seemingly random patterns of aging-related pathologies. Since previous studies in *C. elegans* revealed that as early as by day 1 of adulthood, subpopulations of animals emerge that significantly differ from others in stress resistance and lifespan ^6^, we focused on the concept that specific fluctuating signals during worm development might differentially affect processes that determine and hence individualize lifespan. We decided to explore the idea that reactive oxygen species (ROS), which are well-known signaling molecules and effectors of redox-sensitive processes in cells ^7–9^, might unexpectedly serve as early lifespan determining modulators in *C. elegans*^10^. This hypothesis was validated by a series of independent *C. elegans* studies, which showed i) that significant lifespan extension occurs upon pharmacologically increased ROS generation in the young adult^11^ as well as during development^12^, ii) that exposure of nematodes to non-lethal concentrations of ROS leads to increased stress resistance and longevity^13–15^, a phenomenon termed mitochondrial hormesis or mitohormesis (i.e., health-beneficial effects elicited by exposure of organisms to low concentrations of mitochondria-derived stressors, namely ROS)^11,16^ and iii) that larvae of a synchronized wild-type population exhibit large interindividual variations in endogenous ROS levels ^17^.

## Redox states during development differentially affect redox states in adults

To investigate if and how developmental ROS levels and variations affect *C. elegans* later in life, we decided to sort *C. elegans* L2 larvae according to their endogenous ROS levels and follow them over time. For these experiments, we used wild-type N2 worms that ubiquitously express the integrated redox sensing protein Grx1-roGFP2 ^18^. This cytosolic redox sensor faithfully responds to the cellular ratio of oxidized and reduced glutathione (GSSG:GSH), which is directly affected by a range of different endogenous oxidants, including peroxide ^19^. Consistent with our previous studies in which we used a peroxide-sensor-expressing *C. elegans* strain ^17^, fluorescence microscopy measurements of the GSSG:GSH ratios in live worms revealed a significantly more oxidizing redox environment and substantially higher inter-individual differences in L2 larvae compared to young adults. Coinciding with reaching reproductive activity, the endogenous redox environment of young adult worms appeared to be maximally reduced and showed little inter-individual differences (Extended Data Fig. 1a). As the worms aged, however, the average redox state became more oxidizing again and inter-individual redox differences reemerged.

To be able to sort individuals of a synchronized population according to their respective redox states during early larval development, and obtain sufficient quantities of live animals with similar redox states for follow-up studies, we specifically reconfigured a large particle BioSorter. This reconfiguration was made necessary because the read-out of the Grx1-roGFP2 is ratiometric and demands that each worm is measured by two lasers in series (Extended Data Fig. 1b). Analysis of about 16,000 age-synchronized Grx1-roGFP2-expressing L2 larvae in the BioSorter confirmed our microscopic data and showed that the GSSG/GSH ratio varies widely among individuals (Fig. 1a). We subsequently sorted and binned worms with redox states between 2 and 3 standard deviations above (from hereon: oxidized subpopulation) or below (from hereon: reduced subpopulation) the mean population (Extended Data Fig. 1c and Fig. 1a). We used fluorescence microscopy to confirm that the redox states of individuals sorted into oxidized and reduced subpopulations are indeed significantly different (Fig. 1b and Supplementary Fig. 1). We ascertained that worms in the two subpopulations did not differ significantly in size (Extended Data Fig. 2a) or reproductive activity (Extended Data Fig. 2b), making redox-mediated differences in worm development unlikely. Furthermore, we investigated their mitochondrial respiratory chain function, including basal oxygen consumption rate (OCR), maximal respiratory capacity, and spare respiratory capacity as well as analyzed the basal extracellular acidification rate (ECAR) as proxy for baseline glycolytic flux, but found no significant differences between worms of the two subpopulations (Extended Data Fig. 2c-f).

**Figure 1.**
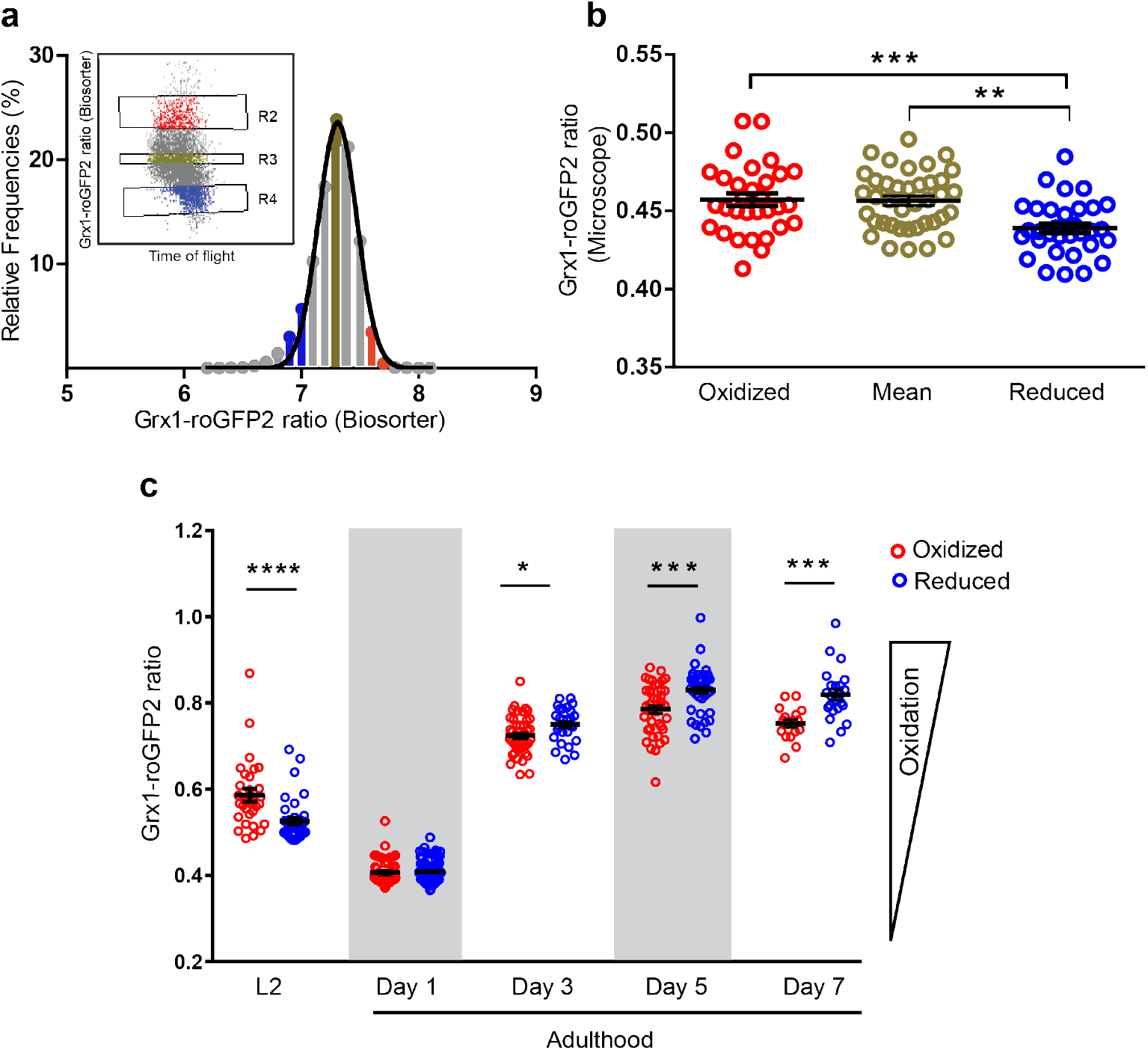
Analysis of the endogenous redox state in an age-synchronized population of *C. elegans*. (a) Distribution of Grx1-roGFP2 ratios of a L2-staged N2*jrIs2[Prpl-17::Grx1-roGFP2]* population according to the BioSorter. L2 worms with Grx1-roGFP2 ratios between 2 and 3 standard deviations (bars) above (insert: R2, red, oxidized) or below (insert: R4, blue, reduced) the mean were sorted based on the peak intensity of the 405 nm/488 nm signals (Extended Data Fig. 1b). L2 worms with average Grx1-roGFP2 ratios were sorted as reference (insert: R3, green, mean). For insert in (1a), axes have been redrawn to scale for clarity. N = 15599 animals. (b) Microscopic analysis of the Grx1-roGFP2 ratio of individual L2 worms (symbol) previously sorted into oxidized (red), mean (green) and reduced (blue) subpopulations. **, *p* <0.01; ***, *p* < 0.001 (one-way ANOVA with Tukey correction). Experiments were performed a minimum of 5 times (see Supplemental Figure 1 for individual results) and a representative graph is shown. (c) Longitudinal analysis of the redox state. Worms were sorted at the L2 larval stage into reduced and oxidized subpopulations, and subsequently cultivated at 20 °C. The Grx1-roGFP2 ratio of animals (symbol) in each subpopulation was determined microscopically at the indicated time points. N = 3 experiments, 17-86 animals per condition. *, *p* < 0.05; ***, *p* < 0.001; ****, *p* < 0.0001 (Mann-Whitney U test). Data in (b) and (c) represent mean ± SEM.

Subsequent redox analysis of worms after their sorting into subpopulations at the L2 larval stage showed that all worms became similarly reduced in young adulthood with no apparent interindividual variation in their redox states (Fig. 1c). However, as the animals matured and the endogenous redox states started to get more oxidizing again, the two populations diverged but now in opposite redox directions (Fig. 1c). At day 7 of adulthood, animals originally sorted into the oxidized subpopulation were more reduced than animals that were originally binned into the reduced subpopulation. At this point, we do not know what event(s) trigger the transient increase in GSSG:GSH ratios during early development, or what mechanisms are responsible for the observed switch in endogenous redox states during adulthood. However, our observations clearly demonstrate that a synchronized, isogenic population of young *C. elegans* larvae contains subpopulations of worms with redox environments that imprint information, which may become relevant later in life.

## Redox state during development affects stress resistance and longevity

To investigate the potential downstream effects of the observed variations in developmental redox levels, we compared stress resistance and lifespan of animals that had been sorted into oxidized and reduced subpopulations at the L2 larval stage. Compared to individuals in the reduced subpopulation, we found that animals in the oxidized subpopulation were significantly more heat shock resistant (Fig. 2a), about 30% longer-lived after the initial heat shock treatment (Fig. 2b), and substantially longer-lived when grown in the presence of oxidants, such as paraquat (Fig. 2c) or juglone (Extended Data Table 1). Moreover, worms sorted into the oxidized subpopulation displayed an up to 18% increase in median lifespan and a 1 - 4 day increase in maximal lifespan (Fig. 2d; Extended Data Table 2 and Supplementary Fig. 2). These results showed a clear correlation between increased GSSG:GSH ratios, increased stress resistance and extended lifespan in a subpopulation of synchronized, age-matched worms experience a naturally occurring hormesis event during early development.

**Figure 2.**
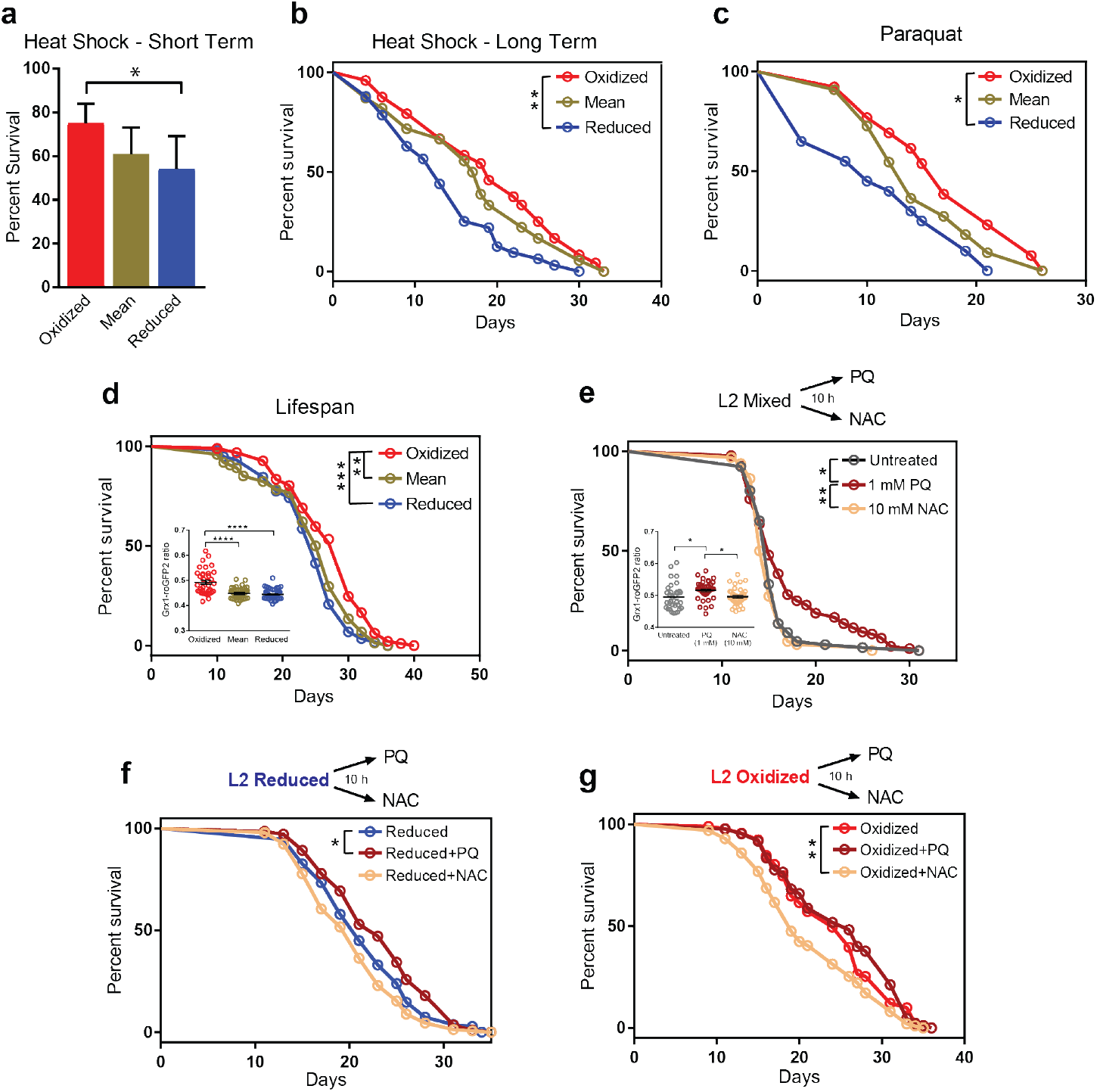
Oxidized L2 subpopulations show increased stress survival and longer lifespan. All experiments were performed with N2*jrIs2[Prpl-17::Grx1-roGFP2]* animals. (a) Survival of oxidized, mean and reduced subpopulations 20 hours after heat shock. N = 5 experiments, 204-331 animals per condition. *, *p*<0.05 (one-way ANOVA with Tukey correction). Data represent mean ± SEM. (b) Representative survival curves of oxidized, mean and reduced subpopulations that survived the heat shock treatment. *, *p*<0.05 (log-rank test). (c) Representative survival curves of oxidized, mean and reduced subpopulations in the presence of 2 mM PQ. *, *p*<0.05 (log-rank test). (d) Representative survival curves of oxidized, mean and reduced subpopulations. **, *p*<0.01 (log-rank test). (Insert) The Grx1-roGFP2 ratio of individual worms (symbol) after sorting is shown. ****, *p* < 0.0001 (one-way ANOVA with Tukey correction). (e) Representative survival curves of a non-sorted (mixed) population of worms treated at the L2 stage with either nothing (gray), 1 mM PQ (red) or 10 mM NAC (yellow) for 10 hours. *, *p*<0.05 (log rank test). (Insert) The Grx1-roGFP2 ratio of individual worms (symbol) after treatment is shown. *, *p*<0.05 (one-way ANOVA with Tukey correction). Data in inserts represent mean ± SEM. (f) Representative survival curves of a reduced subpopulation of L2 worms after a 10 h-treatment with either nothing (blue), 1 mM PQ (red) or 10 mM NAC (yellow). *, *p*<0.05 (log rank test) (g) Representative survival curves of an oxidized subpopulation of L2 worms after a 10 h-treatment with either nothing (red), 1 mM PQ (dark red) or 10 mM NAC (yellow). **, *p*<0.01 (log rank test). The specific sorting events, number of individuals and statistics obtained for each of the data sets shown in panels (b)-(g) can be found in Ext. Data Tables 1–3.

To test whether a drug-induced alteration of the cellular GSSG:GSH ratio specifically during the L2 larval stage is sufficient to affect the lifespan of the entire population, we exposed an unsorted (mixed) population of L2 worms to either no treatment, 1 mM of the oxidant paraquat (PQ) or 10 mM of the reductant N-acetyl cysteine (NAC) for 10 hours. After the treatment, we placed the worms onto NGM plates, and assessed their lifespan. As shown in Fig. 2e, a 10 h treatment of L2 larvae with paraquat was sufficient to significantly extend the maximum lifespan of the whole population. No effect on the population was observed when we treated the worms with N-acetyl cysteine. These results were fully consistent with Grx1-roGFP2 measurements, which showed that treatment with paraquat caused a pronounced shift in the entire population towards a more oxidizing cellular environment, whereas the 10 h N-acetyl cysteine treatment had no detectable effect on the redox state of the population as a whole (Fig. 2e, insert). When we conducted the same experiments with sorted subpopulations, we found that N-acetyl cysteine treatment reduced the lifespan in animals of the oxidized but not of the already reduced subpopulations (Fig. 2f and Extended Data Table 3), whereas paraquat treatment increased the lifespan of animals in the reduced but not in the oxidized subpopulation (Fig. 2g and Extended Data Table 3). These results provide evidence that transient changes in the redox-environment during early development are sufficient to positively affect the lifespan of *C. elegans*. Within a population, however, it appears that only a small subset of worms encounter levels of endogenous ROS during development that are sufficiently high to trigger these beneficial long-term effects.

## Global H3K4me3 levels are redox-sensitive epigenetic marks

To gain insights into the mechanism(s) by which a transient increase in the cellular redox state during development causes an increase in stress resistance and lifespan, we first conducted quantitative RT-PCR, testing for mRNA levels of commonly assessed heat shock genes (*i.e*., the heat stress-induced transcription factor *hsf-1;* the ortholog of human Hsp70, *hsp-1;* and the small heat shock protein *hsp-16.2*) (Fig. 3a) and oxidative stress-related genes (e.g., superoxide dismutases *sod-1*, *2*, *3*; catalases *ctl-1* and *ctl-2*) (Extended Data Fig. 3a). Unexpectedly, we did not detect any significant differences in the steady-state expression levels of these stress-related genes in worms of the two subpopulations. However, upon exposure to heat shock conditions, we discovered that animals of the oxidized subpopulation showed a significantly increased capacity to upregulate heat shock gene expression compared to animals of the reduced subpopulation (Fig. 3a). The mRNA levels of HSP-16.2, for instance, increased by over 700-fold in worms of the oxidized subpopulation compared to less than 200-fold in worms of the reduced subpopulation (Fig. 3a). Transcript levels of the heat shock factor HSF-1 before and after heat shock were not significantly different between the two subpopulations (Fig. 3a), suggesting that the transcriptional stimulation in animals of the oxidized subpopulation is either due to specific changes in HSF-1 activity, its subcellular localization and/or changes in heat shock promoter accessibility ^20,21^.

**Figure 3.**
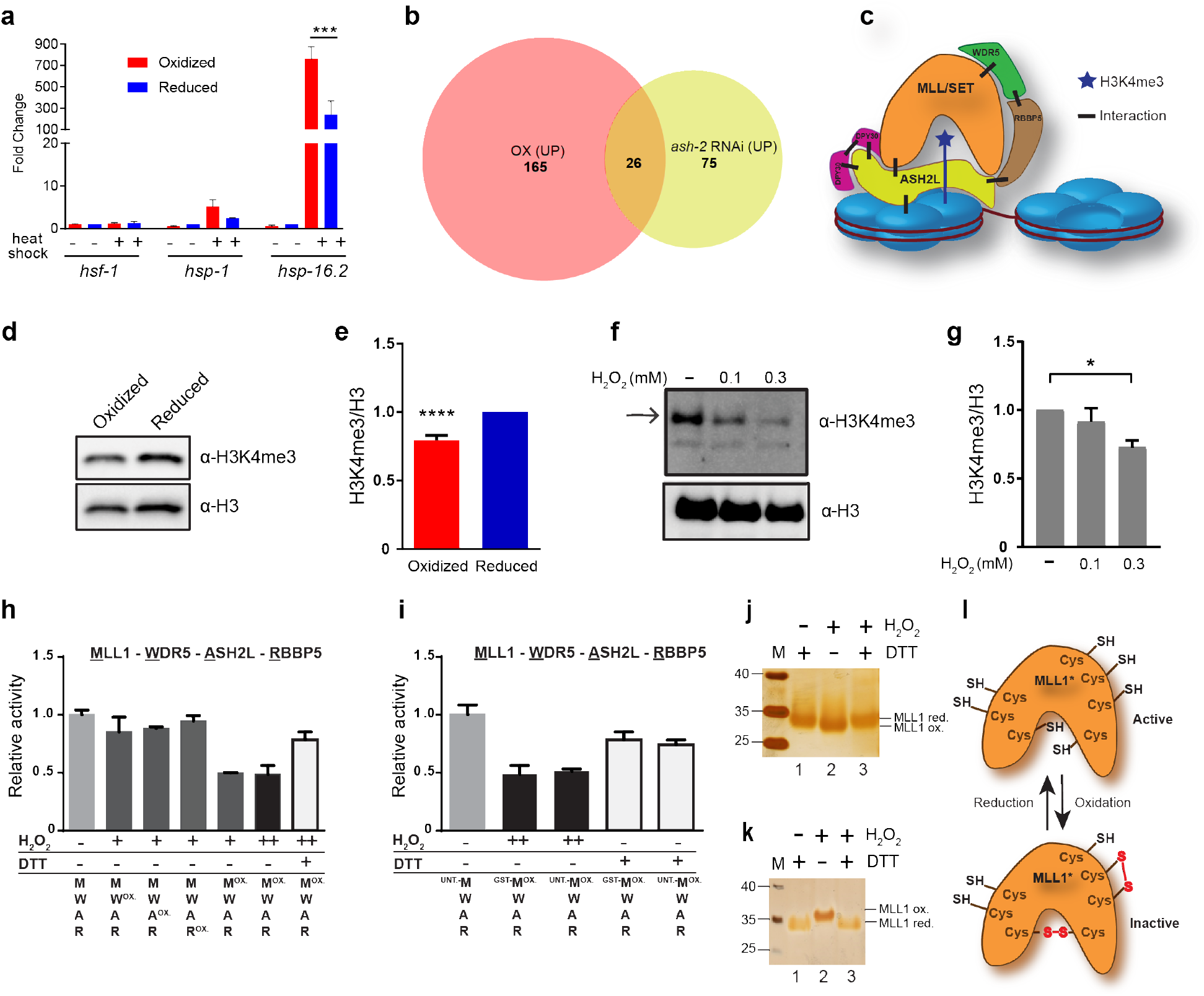
Oxidizing conditions reduce global H3K4me3 levels in *C. elegans* and HeLa cells. (a) Transcript levels of selected heat shock genes before and after heat shock treatment of oxidized and reduced subpopulations at the L2 stage, as assessed by qRT-PCR, using *cdc-42* as a reference gene. N = 3 experiments ****, *p*<0.0001 (one-way ANOVA with Tukey correction). The expression date of the oxidized subpopulation were normalized to the reduced subpopulation, and represent mean ± SEM. (b) Venn diagram of differentially expressed genes (DEGs) upregulated in the oxidized L2 subpopulation and in *ash-2* RNAi knockdown worms ^22^ (for the complete list of overlapping DEGs, see Supplementary Table 1). (c) Schematic representation of the H3K4 methylation complex (HMC), also referred to as KMT2 (class 2 lysine methyltransferase). (d) Global H3K4me3 levels in the oxidized and reduced subpopulations of L2 worms as assessed by western blot. A representative blot is shown. (e) Quantification of global H3K4me3 levels in oxidized and reduced subpopulations at the L2 stage. N = 7 experiments. ****, *p*<0.0001 (unpaired *t*-test). Data were normalized to the reduced subpopulation and represent mean ± SEM. (f) Global H3K4me3 levels in HeLa cells before and after H_2_O_2_ treatment as assessed by western blot. A representative blot is shown. (g) Quantification of global H3K4me3 levels in HeLa cells before and after H_2_O_2_ treatment. N = 3. *, *p*<0.05 (one-way ANOVA with Dunnett correction). Data were normalized to the untreated control and represent mean ± SEM. (h, i) *In vitro* histone methyltransferase assays of core HMC members, consisting of purified GST-WDR5 (WDR5), GST-ASH2L (ASH2L), GST-RBBP5 (RBBP5) and either GST-MLL1_SET_ (h) or MLL1_SET_ (labeled with ^UNT^-M) (i). Superscript OX indicates that the protein was pre-treated with either 1 mM (+) or 2 mM (++) H_2_O_2_ for 30 min prior to the activity assay. DTT was added after the H_2_O_2_ treatment. N = 3. Data represent mean ± SEM. (j, k) MLL1_SET_ treated with either 2 mM DTT, 2 mM H_2_O_2_ or 2 mM H_2_O_2_ followed by 4 mM DTT-treatment was analyzed. (j) All reduced protein thiols present in either MLL1_SET_ preparation were labeled with the 500 Da thiol-reactive AMS, causing a 500-Da mass *de*crease per oxidized thiol detectable on reducing SDS-PAGE. (k) For the reverse thiol trapping, all reduced thiols were first labeled with N-ethylmaleimide (NEM), followed by the reduction of all oxidized protein thiols and their subsequent labeling with AMS, causing a 500-Da mass *in*crease per oxidized thiol detectable on non-reducing SDS-PAGE. (l) Schematic representation of the redox sensitivity of the MLL1_SET_. Data in (a), (e), (g), (h) and (i) represent mean ± SEM. For blot and gel source images, see Supplementary Fig. 3.

RNA-seq analysis from four independent large-scale sorting experiments demonstrated clear differences in gene expression between the two subpopulations (Extended Data Fig. 3b) but did not provide any obvious connection between the 191 up-regulated or 136 down-regulated genes in the oxidized subpopulation to previously investigated sets of stress or longevity-related genes (Extended Data Fig. 3c and Supplementary Table 1). To our surprise, however, we identified a substantial overlap between genes that were upregulated in the oxidized subpopulation with genes upregulated in worms lacking the Absent Small Homeodisc protein ASH-2^22^ (*P* = 1.7×10^−22^, hypergeometric probability) (Fig. 3b and Supplementary Table 1). Specifically, we found 26 of the upregulated genes among the 102 genes upregulated in *ash-2* knockdown worms (expected overlap if there was no correlation is >1). ASH-2 is a component of the highly conserved multiprotein histone methylation complex called COMPASS (Fig. 3c), which, together with a member of the SET-1/MLL (mixed lineage leukemia) histone methyltransferase family (SET-2 in *C. elegans*) and WDR-5 is responsible for the trimethylation of lysine 4 in histone H3 (H3K4me3) in organisms ranging from yeast to humans ^23,24^. H3K4me3 is primarily found at transcriptional start sites (TSS), where the modification is thought to mark and maintain transcriptionally active genes ^25^. Intriguingly, recent studies in *C. elegans* revealed that TSS-associated H3K4me3 marks are set during early development (at or before L3 larval stages), and remain stable over the lifespan of the organism ^19^. Indeed, analysis of published ChIP-data revealed that about 25% of the differentially expressed genes (DEGs) that we identified in the oxidized subpopulation of worms associate with H3K4me3 marks that appear to be set specifically during development (Extended Data Fig. 4a, b). Moreover, comparison with recently conducted gene expression studies in worms lacking the H3K4me3 readers SET-9 or SET-26 revealed highly significant overlaps for both up- and down regulated genes in the oxidized subpopulation with genes differently expressed in *set-9* or *set-26* mutant backgrounds, very similar to the overlaps previously observed with *ash-2* knockdown strains (Extended Data Fig. 4c, d)^26^. Based on this realization that the oxidized subpopulation of L2 worms had gene expression changes consistent with reduced H3K4me3 marks, we tested the two subpopulations for global H3K4me3 abundancy. These experiments specifically showed a > 25% reduction in global H3K4me3 levels in the worms of the oxidized subpopulation compared to worms of the reduced subpopulation (Fig. 3d and e). These results strongly suggested that we have discovered a connection between endogenous ROS levels, H3K4 trimethylation levels, and gene regulation.

## SET1/MLL histone methyltransferases are redox-sensitive proteins

RNA-seq analysis of the oxidized and reduced subpopulation of worms did not reveal any transcriptional changes in components of the H3K4me3 complex that would explain the decrease in H3K4me3 levels (Supplementary Table 1). This result suggested that the activity of one or more member(s) of the H3K4me3 complex might be post-translationally affected by the cellular redox environment. To directly test whether the H3K4me3 machinery is sensitive to ROS, we attempted to purify the *C. elegans* proteins for *in vitro* methylation assays. This, however, turned out to be impossible due to their instability. As an alternative approach, we therefore considered the use of the mammalian H3K4me3 machinery. The components of the COMPASS family are all highly conserved and have been successfully purified and previously used in H3K4 methylation assays *in vitro* ^23,27,28^. To first ascertain that H3K4 trimethylation is a redox sensitive process in mammalian cells as well, we subjected HeLa cells to non-lethal H_2_O_2_ treatment, and investigated the global H3K4me3 levels. Consistent with our findings in *C. elegans*, we observed a > 30% global decrease in H3K4me3 levels within less than 30 min of peroxide treatment (Fig. 3f and 3g), without any detectable changes in the steady-state levels of two of the main COMPASS components (Extended Data Fig. 5a, b).

We then purified the components of the minimal mammalian H3K4me3 machinery that have been shown to methylate H3K4 in *vitro*; the multiple leukemia lineage MLL1 methyltransferase SET-domain (MLL1_SET_), which plays the central role in SET1/MLL proteins by catalyzing the lysine-directed histone methylation^29^, ASH2L (ASH-2 in *C. elegans*), WDR5 (WDR-5.1 in *C. elegans*) and RBBP5 (RBBP-5 in *C. elegans*). We individually treated the four proteins with peroxide for 30 min, removed the oxidant, combined the proteins and tested for the *in vitro* histone methylation activity of the complex. Notably, only one protein appeared to be reproducibly peroxide-sensitive; the SET-domain of MLL1 (Fig. 3h, i and Extended Data Fig. 5c). Importantly, incubation of the peroxide-inactivated SET-domain with the thiol-reducing agent dithiothreitol (DTT) restored the original activity, strongly suggesting that thiol oxidation is responsible for the observed inactivation (Fig. 3h, i and Extended Data Fig. 5c). Subsequent analysis of the SET-domain of other SET1/MLL-family members ^30–32^, including SET1A, the most closely related human homologue of *C. elegans* SET-2 ^33^, SET1B as well as a version of MLL1 that lacks the cysteine-containing GST-tag used for purification (Fig. 3i), confirmed that the specific and reversible sensitivity towards peroxide treatment is a universal feature for the SET1/MLL family (Extended Data Fig. 5d and e). The MLL1_SET_ domain contains seven cysteines, five of which are highly conserved and four of which have been shown to be involved in the coordination of a zinc ion^34,35^ (Extended Data Fig. 6). Since cysteine-containing zinc centers are potentially redox-sensitive units in proteins, we first compared the migration behavior of oxidized and reduced MLL1_SET_ on SDS-PAGE. Indeed, we found that about 50% of the oxidized protein migrated reproducibly faster than the reduced MLL1_SET_, indicative of the formation of one or more intramolecular disulfide bonds (Extended Data Fig. 5f). This result was consistent with the about 50% reduction in enzyme activity that we had observed in our methylation assays (Fig. 3h, i). Increasing the incubation temperature of the peroxide treatment shifted all of the MLL1_SET_ protein into the faster migrating conformation, indicating complete oxidation of the redox-sensitive cysteines (Extended Data Fig. 5f). To determine how many cysteines get oxidized upon peroxide treatment, we conducted direct and reverse thiol trapping on oxidized and reduced MLL1_SET_ using the 500-Da thiol-reactive compound 4-acetamido-4’maleimidylstilbene-2,2’-disulfonic acid (AMS), followed by SDS PAGE^36^ (Fig. 3j, k). Analysis of the migration behavior of thiol-trapped oxidized versus reduced MLL1_SET_ was consistent with the conclusion that peroxide-treatment leads to the formation of two intramolecular disulfide bonds. Subsequent mass spectrometric analysis of *in vitro* modified cysteines confirmed these results and revealed that at least four of the five absolutely conserved cysteines are highly oxidation-sensitive (Fig. 3l, Extended Data Fig. 5g). To our knowledge, this makes H3K4me3 the first histone methylation mark, and MLL1 the first histone methyltransferase known to be regulated by the redox environment of the environment.

## Decrease in H3K4me3 increases heat shock gene expression and stress resistance

Our studies raised the intriguing possibility that the redox-mediated inactivation of the lone SET-1 homologue in the oxidized subpopulation of *C. elegans* larvae (i.e., SET-2) leads to a reduction in global H3K4me3 levels, which, in turn, causes increased stress resistance and longevity. This theory was supported by recent studies, which showed that deleting or knocking down components of the H3K4me3-machinery in *C. elegans* cause a long-lived phenotype^22^. Indeed, when we analyzed *C. elegans* mutants with reduced levels of H3K4me3 due to the deletion of *set-2* or the knockdown of *ash-2* (Extended Data Fig. 7a and b), we observed a significant increase in heat stress resistance compared to the respective control strains as well as a strain deficient in the H3K4me3-demethylase RBR-2 (Fig. 4a and b). In addition, and similar to the results obtained with the oxidized L2 subpopulation, we found that H3K4me3-deficient worms showed a substantially increased transcriptional response upon heat shock treatment (Extended Data Fig. 8a and b). Although slightly counterintuitive since H3K4me3 is considered an activating mark, these results are fully consistent with recent studies in yeast, which showed that reduction of H3K4me3 levels does not significantly alter steady state gene expression levels but causes substantially more robust gene expression changes upon stress treatment ^37^. We obtained very similar results in HeLa cells, where we found that heat-treated ASH2L-depleted cells (Extended Data Fig. 7c) were significantly more stress resistant (Fig. 4c) and responded more robustly with heat shock gene expression than control RNAi cells (Fig. 4d). These results strongly suggested that the downstream effects of H3Kme3 depletion on stress resistance and longevity are conserved events.

**Figure 4.**
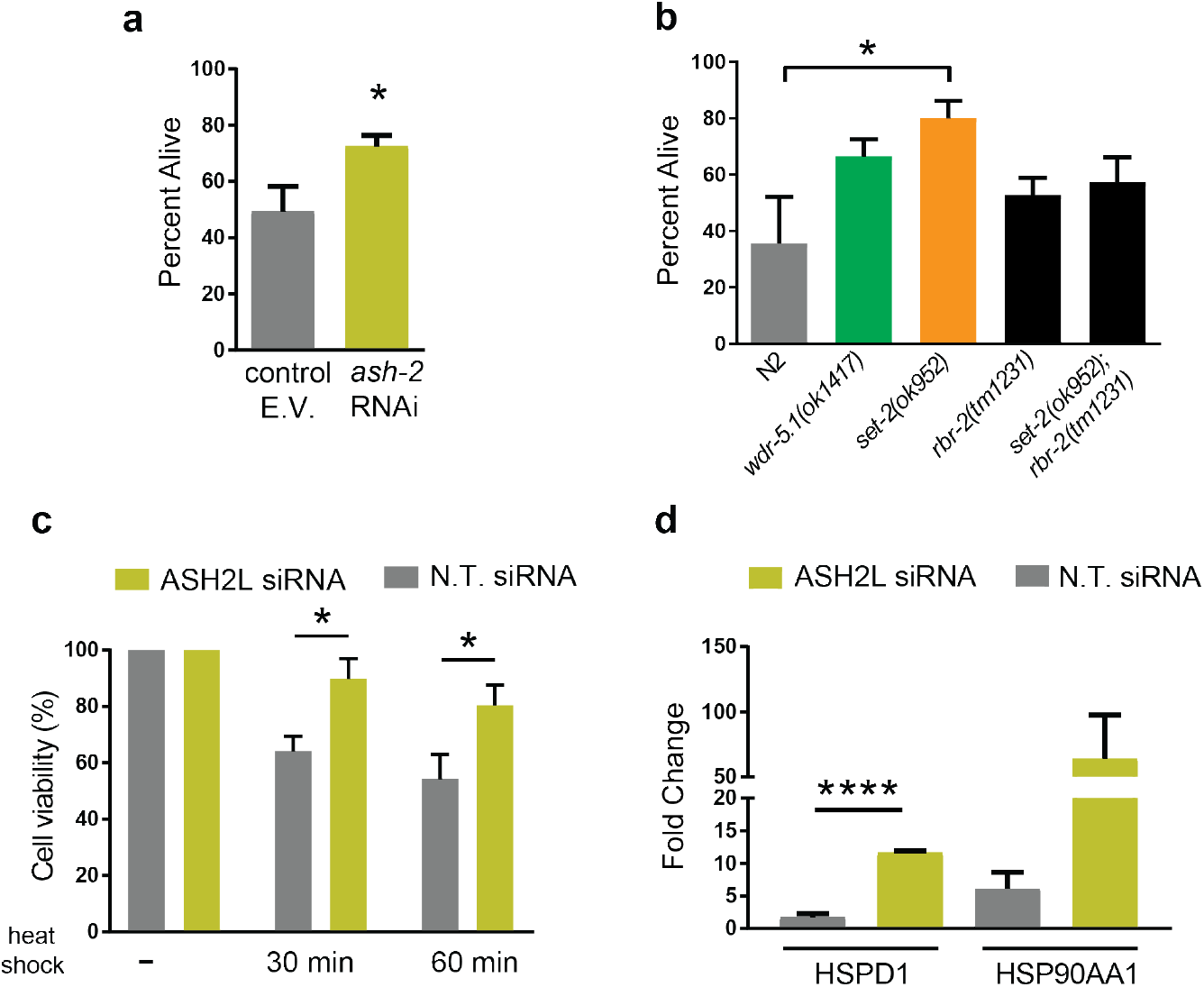
Downregulation of global H3K4me3 levels increases stress resistance. (a) Survival of N2*jrIs2*[P*rpl-17*::Grx1-roGFP2] worms treated with *ash-2* RNAi, measured 24 hours after heat shock. N = 5 experiments. *, *p*<0.05 (unpaired *t*-test). E.V. empty vector. (b) Survival of H3K4me3 deficient mutants, measured 48 hours after heat shock. N = 3 experiments. *, *p*<0.05, (one-way ANOVA with Dunnett correction). (c) Viability of ASH2L siRNA treated HeLa cells upon heat shock. N = 5 experiments. *, *p*<0.05 (two-way ANOVA with Tukey correction). (d) Transcript levels of selected heat shock response genes after heat shock treatment of ASH2L siRNA treated HeLa cells, as assessed by qRT-PCR, using spiked-in luciferase as a reference gene. Data were normalized to the respective non-heat shock control. N.T., nontargeting. ****, *p*<0.0001 (unpaired *t*-test). All data represent mean ± SEM.

To finally test whether down-regulation of H3K4me3 levels is sufficient to increase heat shock resistance and lifespan in the oxidized subpopulation, we generated *ash-2* or *set-2* RNAi knockdown worms expressing the Grx1-roGFP2 sensor protein. As before, we monitored and sorted the synchronized population of L2 larvae into oxidized and reduced subpopulations (Fig. 5a). We did not detect any significant differences in the relative distribution or range of GSSG:GSH ratios between the *ash2* or *set-2* RNAi worm populations and the control RNAi worms (Extended Data Fig. 8c). These results were consistent with the model that an increase in the GSSG-GSH ratio serves at the initial trigger that affects relative H3K4m3 levels in a subpopulation of L2 larvae and sets all subsequent events in motion. More importantly, however, we found that the sorted subpopulations of *ash-2* and *set-2* RNAi worms no longer exhibited any difference in heat shock sensitivity (Fig. 5b and c) or lifespan (Fig. 5d and e; Extended Data Tables 2 and 4). Similarly, no life-prolonging effect was observed when we treated *ash-2* or *set-2*-RNAi-worms with PQ for 10h-treatment at the L2 larval state (Extended Data Fig. 8d and Extended Data Table 4). These results implied that down-regulation of H3K4me3 levels is both necessary and sufficient to increase heat shock resistance and lifespan in the oxidized subpopulation of worms (Fig. 5f).

**Figure 5.**
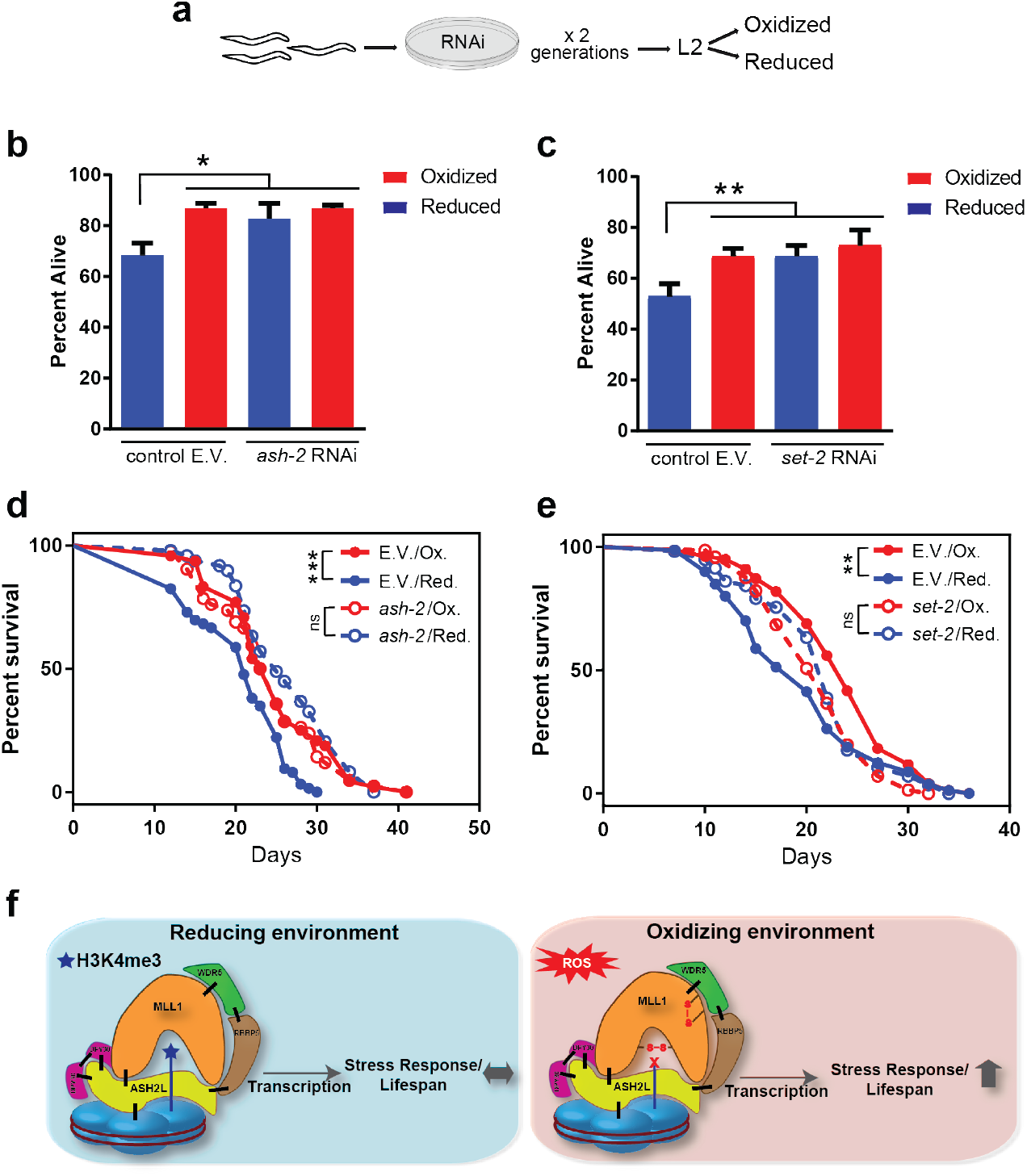
An intrinsically oxidizing environment confers increased stress resistance via downregulation of global H3K4me3 levels. (a) Scheme (top) of heat shock resistance assessment of oxidized and reduced worm subpopulations treated with *ash-2* RNAi. Survival of N2*jrIs2*[P*rpl-17*::Grx1-roGFP2] worms treated with *ash-2* (b) or *set-2* (c) RNAi, sorted into reduced and oxidized subpopulations at the L2 stage and measured 24 hours after heat shock. N = 3 (b) and N = 4 (c) experiments. Data were normalized to the respective non-heat shock control. *, *p*<0.05; **, *p*<0.01 (two-way ANOVA with Tukey correction). Data represent mean ± SEM. Representative survival curves of N2*jrIs2*[P*rpl-17*::Grx1-roGFP2] worms treated with *ash-2* (d) or *set-2* (e) RNAi and sorted in reduced and oxidized subpopulations at the L2 stage. **, *p*<0.01; *ns*, not significant (log rank test). For lifespan values and n numbers in (d) and (e), see Extended Data Table 4. (f) Model. An endogenous oxidizing environment during early development promotes increased stress resistance and extended lifespan through a decrease the H3K4-trimethylation events.

## Conclusions

In 2010, Cynthia Kenyon wrote in the journal Nature: “*It is possible that a stochastic event — a metabolic insult or noise in the expression of a regulatory gene — flips an epigenetic switch or sets in motion a chain of events that promotes ageing”* ^38^. Our studies now identified that variations in endogenous ROS during development, potentially caused by locally different growth conditions, contribute to the lifespan variation observed in synchronized populations of *C. elegans*. Animals that accumulate high levels of ROS during development apparently undergo an endogenous hormesis event, which, as previously observed upon exogenous ROS treatment ^13–15^, increases stress resistance and lifespan. This might serve as a bet-hedging strategy to provide subpopulations of worms with improved survival during stress. Our studies uncovered the underlying mechanism of ROS-mediated hormesis by demonstrating that global H3K4me3 levels are redox-regulated, and decrease in response to oxidative stress. Based on the findings that reduction of global H3K4me3 levels increase stress resistance and *C. elegans* lifespan^22^, we now postulate that we have indeed identified one stochastic event, the respective epigenetic switch and the chain of events that are set into motion during early development to increase lifespan.

Recent studies in *C. elegans* demonstrated that H3K4me3 marks within gene bodies are set during adulthood and change with age^19^. In contrast, H3K4me3 that are enriched at transcriptional start sites and thought to provide the memory of actively transcribed genes, are set during development and remain stable throughout the lifespan^19^. This result likely explains how transient redox-mediated changes in H3K4me3 levels during development are sufficient to exert long-lasting effects despite the dramatic changes in the redox environment during adulthood. Our finding that organisms with reduced levels of H3K4me3 (either elicited by an oxidizing redox environment or by genetic means) show an increased transcriptional capacity to respond to stress conditions, such as heat shock (our study) or oxidative stress ^37^, serves to explain their increased stress resistance. However, the extent to which this increase in transcriptional capacity of stress-related genes is linked to the observed lifespan extension remains to be determined. Moreover, while our gene expression studies did not reveal any major enrichment for genes of known longevity pathways, we did observe a significant number of genes involved in lipid metabolism to be downregulated in the oxidized L2-subpopulation (Extended Data Fig. 3c, Supplementary Table 1). It is of note that increased lipid storage and altered lipid signaling have been previously linked to increased lifespans in a variety of different organisms ^39^, and found to play a role in the lifespan extension of worms globally defective in H3K4me3 levels ^40^. Future work is needed to reveal how the transient downregulation of H3K4me3 levels selectively during development can elicit similarly profound life-altering effects. Our ability to change the lifespan of an entire population by a simple 10-hour exposure to reactive oxygen species during development suggests that we have indeed identified a window in time and a mechanism that helps to individualize lifespan in animals. This study will be forming the groundwork for future work in mammals where very early and transient metabolic events in life seem to have equally profound impacts on lifespan ^41^.

## METHODS

### *C. elegans* strains, maintenance and lifespan assays

The following *C. elegans* strains were used in this study: PB020: N2*jrIs2*[P*rpl-17::*Grx-1-roGFP2], N2: Wild-type Bristol isolate, RB1304: *wdr-5.1(ok1417)*, RB1025: *set-2(ok952)*, ZR1: *rbr-2(tm1231)* and ABR9: *set-2(ok952);rbr-2(tm1231)*. If not stated otherwise, worms were cultured at 20°C. Standard procedures were followed for *C. elegans* strain maintenance ^42^. Synchronization was performed using alkaline hypochlorite solution; eggs were allowed to hatch by overnight incubation in M9 medium during gentle shaking. Newly hatched, arrested L1 larvae were transferred onto standard nematode growth medium (NGM) plates seeded with live *E. coli* OP50. Lifespan studies were performed at 20°C in the presence of FuDR. Survival was scored every 2 days, and worms were censored if they crawled off the plate, hatched inside, or lost vulva integrity during reproduction. The first day of adulthood was set as t = 0. Lifespan of unsorted worm populations were performed in a lifespan machine according to ^43^. Survival plots were generated using GraphPad Prism. Lifespan data were analyzed for statistical significance with log-rank (Mantel-Cox) or Gehan-Breslow-Wilcoxon test.

### Reconfiguration of BioSorter for ratiometric sorting

405 and 488 nm lasers were used to excite the Grx1-roGFP2 sensor protein. Since the protein possesses a single emission maximum (~520 nm), the two lasers in the BioSorter (Union Biometrica) were realigned to sequentially illuminate single L2-staged worms as they pass through the flow cell, without emitting overlapping signals. This enabled collection of signals from 405 and 488nm lasers separately, from two photon multipliers tubes (PMTs). As result, data were displayed as two groups of peaks (Extended Data Fig. 1b). Using the partial profiling feature (pp) of the FlowPilot-Pro™ software, we mapped the peaks corresponding to each laser that trace the fluorescent intensity and extinction signals. The extinction signal from the 488 nm laser was used to initially gate worms at the L2 stage larva (R1 gate, see Extended Data Fig.1c). Oxidized, mean and reduced L2 worms were sorted from R2, R3 and R4 gates respectively, based on the peak 405 and 488 fluorescent intensities (insert in Fig. 1a).

### Microscopy

Worms were mounted on objective slides using 4 μl thermoreversible CyGEL (BioStatus; Fisher Scientific) and 2 μl of 50 mM levamisole for immobilization. Fluorescence and DIC images were acquired with an upright microscope equipped with a Photometrics Coolsnap HQ2 cooled CCD camera, a UPlan S-Apo 20x objective (NA 0.75) and a X-CITE^®^ exacte light source equipped with a closed feedback-loop. For Grx1-roGFP2 fluorescence, an external filter wheel was used with excitation filters 420/40x, 500/20x, dual bandpass dichroic T515LPXR and a single emission filter 535/30x. Image analysis was performed in Metamorph (Molecular Devices, Inc) using a custom script. Briefly, an intensity threshold was chosen by the user. Pixels above this threshold constitute regions of interest. Regions with very high signal in any channel (e.g., fluorescent particles) were identified by applying an over-saturation threshold and excluded from regions of interest. Mean ratiometric values (Ex420Em535/Ex500Em535) of regions of interest are calculated after subtraction of background. Acquisition parameters were kept identical across all samples. For body length measurements, worms were measured from the nose to the tail tip and analysis was performed with ImageJ.

### Brood size

L4-staged worms were transferred onto NGM plates and incubated at 15°C. The parental animals were transferred daily to individual NGM plates until the end of the reproductive period. The progeny of each animal was counted at the L2 or L3 stage.

### Cellular respiration

Real-time oxygen consumption rates (OCR) and extra-cellular acidification rate (ECAR) were measured with a Seahorse XF^e^96 Analyzer (Seahorse bioscience Inc., North Billerica, MA, USA) as described^44^. Briefly 100 L2–staged worms were sorted directly into individual wells of 96-well Seahorse utility plates at a final volume of 200 ul of 10% M9. Acute effects of pharmacological inhibitors FCCP (ETC accelerator) and sodium azide (NaN_3_, Complex IV and V inhibitor) were evaluated by injecting them during the run at final concentrations of 20 uM and 40 mM respectively.

### Heat shock treatment

Heat shock was performed on solid OP50-seeded NGM plates wrapped in parafilm and submerged in a pre-heated water bath. For thermotolerance assays, worms were heat-shocked for 45 min at 38°C. Survival was scored after 24 hours and then until the death of the last worm by absence of touch response or pharyngeal pumping. For transcriptional response assays, worms were heat-shocked for 30 min at 35°C. After 1 hour recovery at 20°C, worms were harvested and snap-frozen in liquid nitrogen. All heat shock treatments were applied to worms at the L2 stage.

### Treatments with N-acetyl cysteine (NAC) and paraquat (PQ)

For survival assays, worms were cultivated on solid OP50-seeded NGM plates, supplemented with the indicated concentrations of NAC or PQ. Survival was determined by absence of touch response or pharyngeal pumping. For transient exposure to compounds, worms were transferred into M9-media supplemented with OP50 and the indicated concentrations of NAC or PQ for 10 hours. Worms were harvested, washed three times with M9 and transferred onto regular OP50-seeded NGM plates.

### RNA extraction and real-time qPCR

3,000-5,000 worms at the L2 stage (whole population or after sorting) were grounded in Trizol reagent (Life Technologies) with sea sand and pestle. After filtering of the sand, samples were vigorously shaken with chloroform, allowed to stand for 3 minutes at room temperature, and then centrifuged at 16,000 g at 4°C. The aqueous phase was then collected and RNA was purified using QIAGEN RNeasy RNA extraction columns as per manufacturers’ recommendations. HeLa cells were directly lysed in the culture dish by adding Trizol, as per manufacturers’ recommendations. For RNA isolation, after addition of ethanol, the lysate was loaded onto QIAGEN RNeasy RNA extraction columns. cDNA synthesis was performed using PrimeScript™ 1st strand cDNA Synthesis Kit (Takara) and real-time quantitative PCR was performed using Radiant™ Green Lo-ROX qPCR Kit (Alkali Scientific), as per manufacturers’ recommendations, in a Eppendorf Mastercycler *ep*gradient S realplex^2^ detection system. Relative expression was calculated from Cycle threshold values using the 2^−ΔΔCt^ method and the expression of genes of interest were normalized to housekeeping genes and/or spiked-in luciferase (10 pg/ml Trizol).

Primers used were *cdc-2*: 5’-AGCCATTCTGGCCGCTCTCG-3’ and 5’-GCAACCGCCTTCTCGTTTGGC-3’; *pmp-3*: 5’-TTTGTGTCAATTGGTCATCG-3’ and 5’-CTGTGTCAATGTCGTGAAGG-3’; *panactin*: 5’-TCGGTATGGGACAGAAGGAC-3’ and 5’-CATCCCAGTTGGTGACGATA-3’; *sod-1*: 5’-AAAATGTGGAACCGTGCTG-3’ and 5’-TGAACGTGGAATCCATGAA-3’; *sod-2*: 5’-GATTTGGAGCCTGTAATCAGTC-3’ and 5’-GAAGAGCGATAGCTTCTTTGAC-3’; *sod-3*: 5’-CACTATTAAGCGCGACTTCGG-3’ and 5’-CAATATCCCAACCATCCCCAG-3’; *ctl-2*: 5’-ATCCCAACATGATCTTTGA-3’ and 5’-TGAGATTCTTCACTGGTTG-3’; *prdx-2*: 5’-CGACTCTGTCTTCTCTCAC-3’ and 5’-GAAGATCATTGATGGTGAT-3’; *aak-2*: 5’-AAGTCTGGAGTTGGGAATACG-3’ and 5’-GTATGCACTTCTTTGTGGAACC-3’; *hsf-1*: 5’-TCCGTATAAGAATGCGACTAGG-3’ and 5’-TAGCTTCTGATGTGGTTGAAGG-3’; *hsp-1*: 5’-GGACGTCTTTCCAAGGATGA-3’ and 5’-TCAAGATCTCGTCGACTTG-3’; *hsp-16.2*: 5’-CTGTGAGACGTTGAGATTGATG-3’ and 5’-CTTTACCACTATTTCCGTCCAG-3’; *hsp-6*: 5’-GATAAGATCATCGCTGTCTACG-3’ and 5’-GTGATCGAAGTCTTCTCCTCCG-3’; *ash-2*: 5’-CGATCGAAACACGGAACGA-3’ and 5’-TGCCGGAATCTGCAGTTTTT-3’; *wdr-5.1*: 5’-CCCTGAAACAATACACTGGACACG-3’ and 5’-AACTGGATGACAATCGGAGGC-3’, HSPD1: 5’-TGCTGAGTTTTGAATGAGCAA-3’ and 5’-CAATCTGCTCTCAAATGGACA-3’; Hsp90AA1: 5’-GAAATCTGTAGAACCCAAATTTCAA-3’ and 5’-TCTTTGGATACCTAATGCGACA-3’; Luciferase: 5’-ACGTCTTCCCGACGATGA-3’ and 5’-GTCTTTCCGTGCTCCAAAAC-3’.

### RNA-seq analysis

Total RNA from 4 biological replicates of worms sorted at the L2 stage (extracted as described above) was assessed for quality using the TapeStation (Agilent). Samples were prepared using the Illumina TruSeq Stranded Total RNA Library Prep kit (Illumina). 100 ng of total RNA was rRNA-depleted using Ribo-Gone (Takara Bio USA). The rRNA-depleted RNA was then fragmented and copied into first strand cDNA using reverse transcriptase and random primers. The products were purified and enriched by PCR (15 cycles) to create the final cDNA library. The 3’ prime ends of the cDNA were adenylated and ligated to adapters, including a 6-nt barcode unique for each sample. Final libraries were checked for quality and quantity by TapeStation and qPCR using Kapa’s library quantification kit for Illumina Sequencing platforms (Kapa Biosystems, Wilmington MA). The samples were pooled, clustered on an Illumina cBot and sequenced on one lane of an Illumina HiSeq4000 flow cell, as paired-end 50 nt reads. The quality of the raw reads data for each sample (e.g. low-quality scores, over-represented sequences, inappropriate GC content) was checked using FastQC (version v0.11.3). The Tuxedo Suite software package was used for the computational analysis of the RNA sequencing ^45,46^. Briefly, reads were aligned to the reference genome WS220 using TopHat (version 2.0.13) and Bowtie2 (version 2.2.1.). Cufflinks/CuffDiff (version 2.1.1) was used for expression quantitation, normalization, and differential expression analysis, using reference genome WS220. For this analysis, we used parameter settings: “--multi-read-correct” to adjust expression calculations for reads that map in more than one locus, as well as “--compatible-hits-norm” and “--upper-quartile– norm” for normalization of expression values. Diagnostic plots were generated using the CummeRbund R package. Genes and transcripts were identified as being differentially expressed based on three criteria: test status = “OK”, FDR ≤0.05, and fold change ≥±1.5. The Bioconductor Package GSA was used to perform enrichment test analysis. The algorithm was modified from the original “Gene Set Enrichment Analysis” (GSEA) ^47^, for better power. Genesets were downloaded from sources indicated in Supplementary Table 1. All FDR corrected *p*-values in this result are extremely significant (FDR~0). RNA-sequencing data described in this study have been deposited in the Gene Expression Omnibus (GEO) database ^48^ under accession number GEO: GSE78990.

### Western blot

Standard methods for western blotting were used for the detection of proteins from worm lysates. Briefly, 3,000-5,000 L2-staged worms were collected in 20 μl of M9 buffer and snap frozen in liquid nitrogen. Laemmli loading buffer was added to the worm pellet (1:1 volume) and the samples were boiled for 5 min, separated by SDS-PAGE and transferred to PVDF membranes. Blots were blocked for 1 hour with 5% milk in PBS and probed with anti-H3 (Abcam, ab1791; 1:2,000), anti-H3K4me3 (Abcam, ab8580; 1:1,000), anti-ASH-2 (Abmart, X3-G5EFZ3, 1:1,000;) or anti-β-tubulin (Santa Cruz, sc-5274; 1:2,000) primary antibodies overnight at 4°C. For the extraction of mammalian proteins, HeLa cells were treated with trypsin (Invitrogen, 25200056) washed twice with PBS and collected in lysis buffer (RIPA + 1 mM PMSF + protease inhibitor cocktail + 1 mM EDTA/0.5 ml lysis buffer per 5 x 10^6^ cells). Samples were incubated for 45 min at 4°C with constant agitation. Lysates were spun down (4°C, 20 min, 12,000 rpm) and snap frozen in liquid nitrogen. Laemmli loading buffer was added to the lysates (1:1 volume) and the samples were boiled for 5 min, separated by SDS-PAGE and transferred to PVDF membranes. Blots were blocked for 1 hour with 5% milk in PBS and probed with anti-H3 (Abcam, ab1791; 1:2000), anti-H3K4me3 (Abcam, ab8580; 1:1,000), anti-ASH2L (Bethyl laboratories, polyclonal, A300-489A; 1:1,000), anti-ASH-2 (Abmart, monoclonal, X-G5EFZ3; 1:1000), anti-ASH2L (Bethyl laboratories, polyclonal, A300-489A; 1:1,000), anti-MLL1 (Bethyl laboratories, polyclonal, A300-374A; 1:500) or anti-β-tubulin (Santa Cruz, sc-5274; 1:2,500) primary antibodies overnight at 4°C. HRP conjugated anti-rabbit (ThermoScientific, 31460) and anti-mouse (ThermoScientific, 31430) secondary antibodies were used at 1:5,000 dilution for 1 h at room temperature. Proteins were detected using Clarity ECL Western blotting substrate (Biorad) and signal was captured using a BioRad ChemiDoc Touch imaging system.

### *C. elegans* RNAi

Escherichia coli HT115 (DE3) strains transformed with vectors expressing dsRNA of the genes of interest were obtained from the Ahringer library (a gift from G. Csankovszki), sequence-verified and grown at 37°C as per manufacturer’s recommendations. L1 worms obtained from synchronized populations were placed onto NGM plates containing ampicillin (100mg/ml^−1^) and IPTG (0.4 mM) seeded with the respective bacteria. Worms were cultivated on either RNAi or the empty vector control bacteria for two generations.

### Mammalian cell culture and H_2_O_2_ treatment

HeLa (EM-2-11ht) cells ^49^ (a gift from J. Nandakumar) were cultured in DMEM (Life Technologies, 11995-065), supplemented with 10% Fetal Bovine Serum (Sigma-Aldrich, F4135) and 1% Penicillin-Streptomycin (Gibco, 15140-122) at 5% CO_2_. At 80% confluency, cells were washed with PBS (Life Technologies, 10010023) and treated with HBSS (Life Technologies, 14025-092) supplemented with 0.1 mM or 0.3 mM H_2_O_2_ and incubated at 37 °C for 30 min.

### Mammalian siRNA and heat shock treatment

8,000 cells were transfected with 4.8 pmoles of ASH2L siRNA (Dharmacon, M-019831-01-0005) or non-targeting siRNA (Dharmacon, D-001210-02-05) using Lipofectamine RNAiMax (Invitrogen, 13778-150) in OPTI-MEM I Reduced serum medium (Gibco, 31985-062). For heat shock treatment, cells were washed with PBS 72 hours after siRNA transfection and placed in HBSS. Plates were wrapped with parafilm and submerged in a pre-heated water bath at 43 °C for the indicated time points. Viability was determined using the CellTiter-Glo Kit (Promega) as per manufacturer’s recommendations. Luminescence was monitored on a FLUOstar Omega microplate reader (BMG Labtech).

### Histone methyltransferase activity assays

SET domains of MLL/SET family proteins (MLL1, SET1A, SET1B), RBBP5 (full length), ASH2L (full length), and WDR5 (full length) were purified as previously described ^50^. The purified proteins were diluted to 10 μM and incubated with 1 mM or 2 mM H_2_O_2_ at 4°C for one hour in the buffer 25 mM Tris-HCl, pH 8.0. For DTT recovery, 4 mM DTT was added after H_2_O_2_ treatment and was incubated at 4°C for 60 min. After oxidation, excess H_2_O_2_ was removed by ultrafiltration. Methyltransferase assays were performed using H3 peptides (residues 1-20) with one additional Tyr-residue at C-terminus for accurate quantification of peptides. An enzyme-coupled continuous spectrophotometric assay system was employed to monitor the time course of the reaction^51,52^. This assay system, which monitors the appearance of the cofactor product (SAH) at an absorbance of 515 nm (i.e., OD_515_) contained the following components: 25 mM Tris (pH 8.0), 320 nM AdoHcy nucleosidase, 480 nM adenine deaminase, 40 U/L xanthine oxidase, 20,000 U/L horseradish peroxidase, 4.5 mM 3,5-dicholoro-2-hydroxybenzenesulfonic acid, 0.894 mM 4-aminophena-zone, 40 μM MnCl_2_, 2.25 μM K_4_Fe(CN)_6_·3H_2_O, 200 μM S-Adenosyl-Methionine and 1 μM of the four mammalian proteins that constitute the minimal H3K4-methylating complex. All components were mixed in 30 μl volume in 384-well plate at RT, and the reaction was initiated by adding 400 μM H3 peptide substrate. The OD_515_ was monitored using a Synergy Neo Multi-Mode Reader (Bio-Tek) for 1 hour at 28 °C. The slope of OD_515_ vs. time from the first 20 minutes linear range was converted into reaction rates. The relative activity for each complex without any H_2_O_2_ pretreatment was set to 1. A buffer control was used to determine the baseline.

### *In vitro* protein oxidation and thiol trapping

Purified MLL1_SET_ (15 μM) was treated with either 2 mM DTT or 2 mM H_2_O_2_ for 30 min at 4°C or 30°C. To stop the reaction, the H_2_O_2_-treated samples were mixed with catalase (0.5 mg/ml). To reduce reversible thiol modifications the oxidized sample were treated with 4 mM DTT for 30 min at 30°C. The reduced cysteines were blocked with 20 mM NEM prior to SDS-PAGE analysis. For reverse thiol trapping experiments, the samples were resuspended in a denaturing thiol-trapping buffer (2.3 M urea, 0.2% SDS, 10 mM EDTA, 200 mM Tris-HCl, pH 8.5) supplemented with 20 mM NEM for 30 min at 25°C. Proteins were precipitated with 10% trichloracetic acid (TCA). After centrifugation, the pellets were washed with 10% TCA and 5% TCA and re-dissolved in the denaturing thiol-trapping buffer supplemented with 4 mM DTT to reduce reversible thiol modifications. After 45 min of incubation at 30°C, all new cysteine thiols were labeled with 25 mM AMS for 5 min at 25°C. Proteins were analyzed on SDS-PAGE under non-reducing conditions and visualized using silver staining. For the mass spectrometric analysis of the cysteine-containing peptides, iodoacetamide (IAM) was used instead of AMS to label reversibly oxidized cysteines. After SDS-PAGE under non-reducing conditions and Coomassie staining, protein bands were cut out, trypsin-digested, and analyzed by nano LC-MS/MS (MS Bioworks).

### Statistical analysis

The Prism software package (GraphPad Software 7) and the Microsoft Office 2010 Excel software package (Microsoft Corporation) were used to carry out statistical analyses. Information about *p* values and experiment specifics (n, statistical significance tests) for all figures are provided in their respective figure legends.

---

**Extended Data Figure 1.**
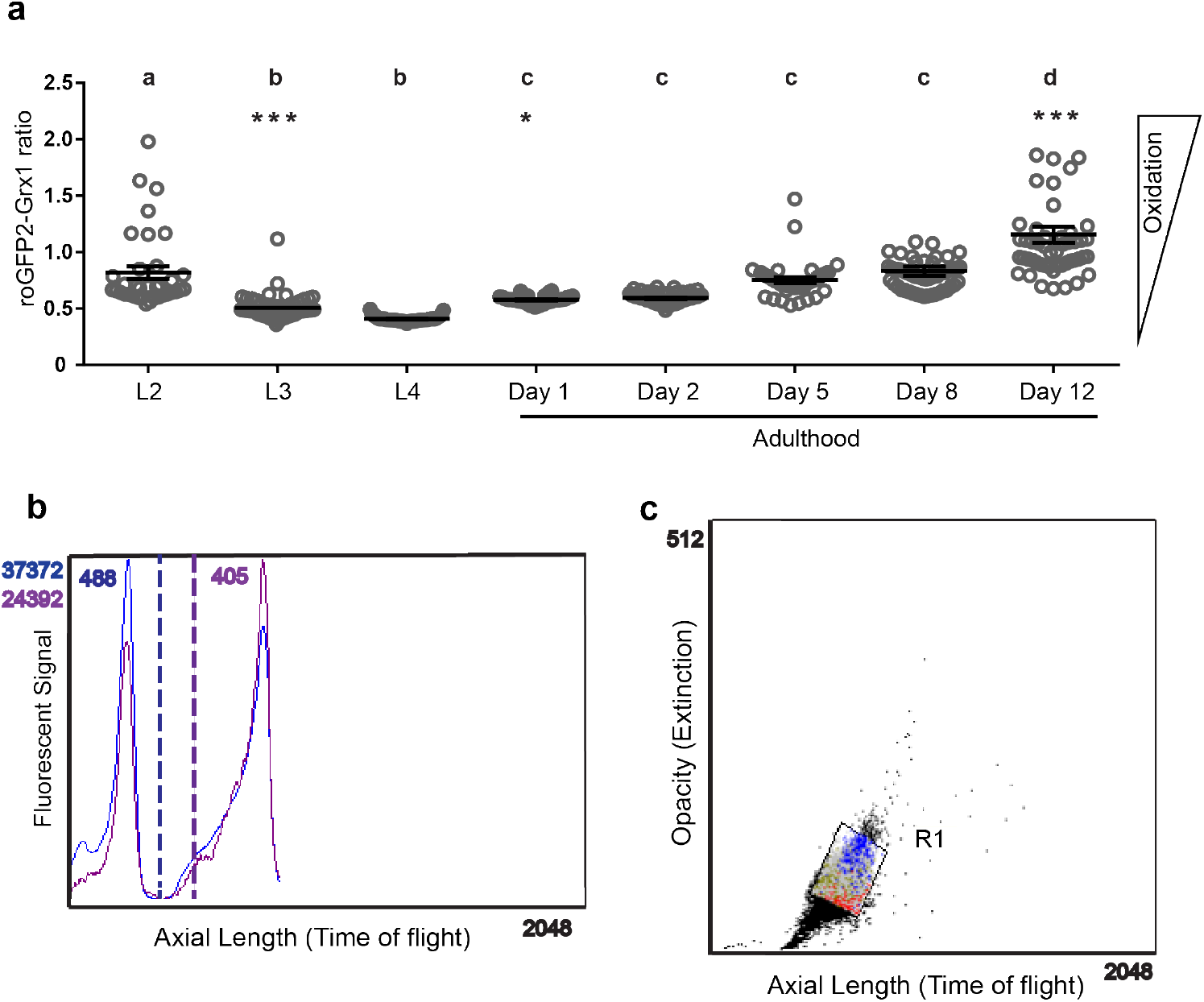
*In vivo* read-out of endogenous redox states at different stages during *C. elegans* lifespan and sorting parameters of oxidized and reduced subpopulations. (a) Microscopic analysis of the Grx1-roGFP2 ratio of individual N2*jrIs2*[*Prpl-17*::Grx1-roGFP2] worms (symbol) cultivated at 15°C and imaged at the indicated time points. Means (bars) of every two sequential time points that are not significantly different from each other (*p*>0.5) share the same letter. Data represent mean ± SEM. *, *p* < 0.05; ***, *p* < 0.001 (one-way ANOVA with Tukey correction). N = 36-54 animals per condition. (b) The roGFP2 ratio (405/488) was calculated using the partial profiling feature (pp) configured to analyze extinction and emission data from each 488 nm and 405 nm lasers that sequentially excited each worm. (c) A population of N2*jrIs2[Prpl-17::Grx1-roGFP2]* at the L2 stage separated based on their opacity (extinction) and length (time of flight) was gated as R1.

**Extended Data Figure 2.**
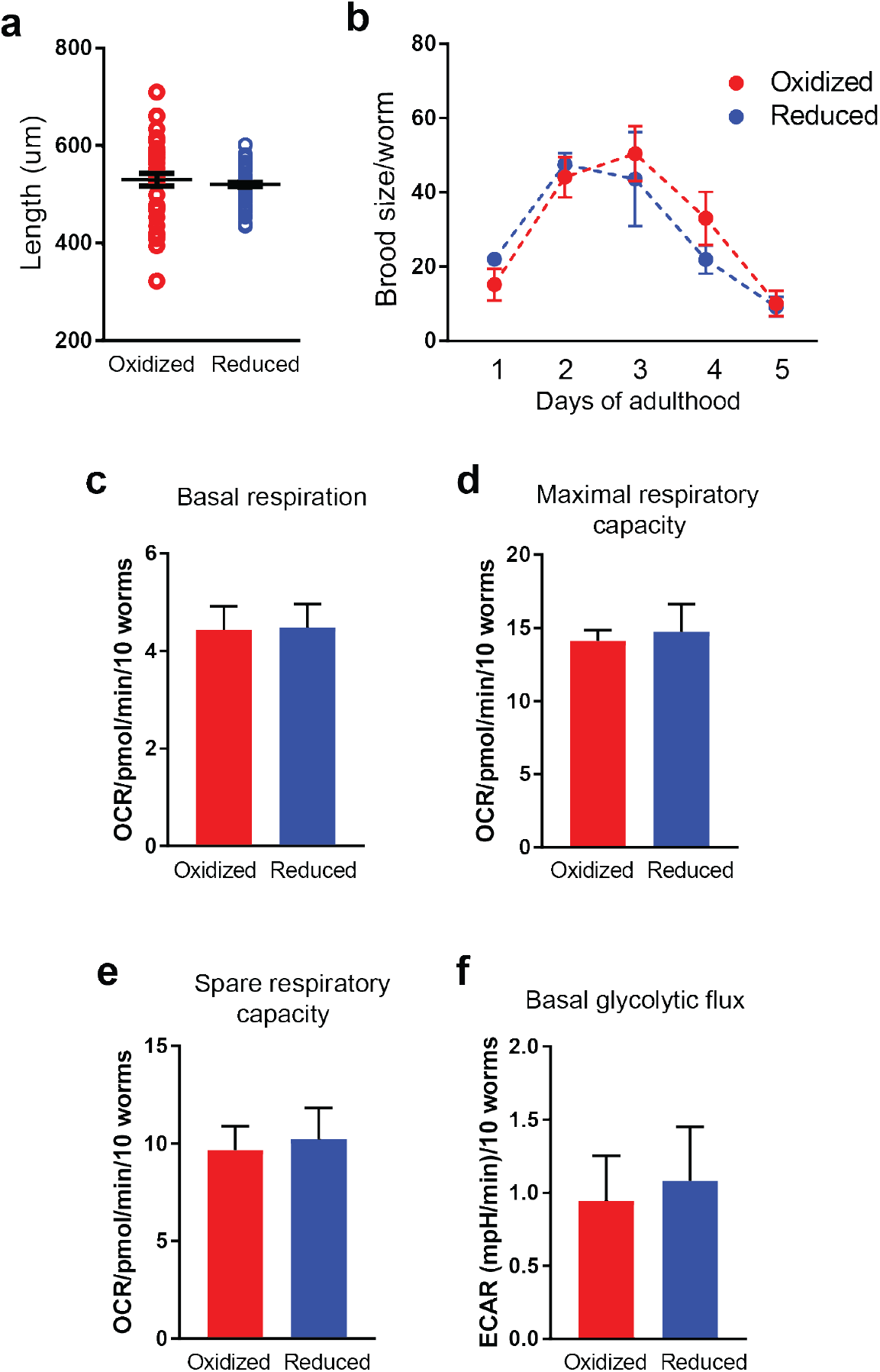
Physiological properties of oxidized and reduced subpopulations at the L2 stage. (a) Length measurements of oxidized and reduced N2*jrIs2[Prpl-17::Grx1-roGFP2]* worms (symbol) from nose to tail tip at the L2 stage. N = 39-49 animals per condition. No significant difference (unpaired *t*-test). (b) Brood size of oxidized and reduced individuals sorted at the L2 stage, measured at the indicated time points. N = 41-125 worms per condition. No significant difference of oxidized versus reduced within a single age (two – way ANOVA). (c) Basal respiration, (d) maximal and (e) spare respiratory capacity and (f) basal rates of flux through glycolysis (ECAR) of oxidized and reduced individuals sorted at the L2 larval stage. N = 3 experiments. No significant difference (unpaired *t*-test). All data represent mean ± SEM.

**Extended Data Figure 3.**
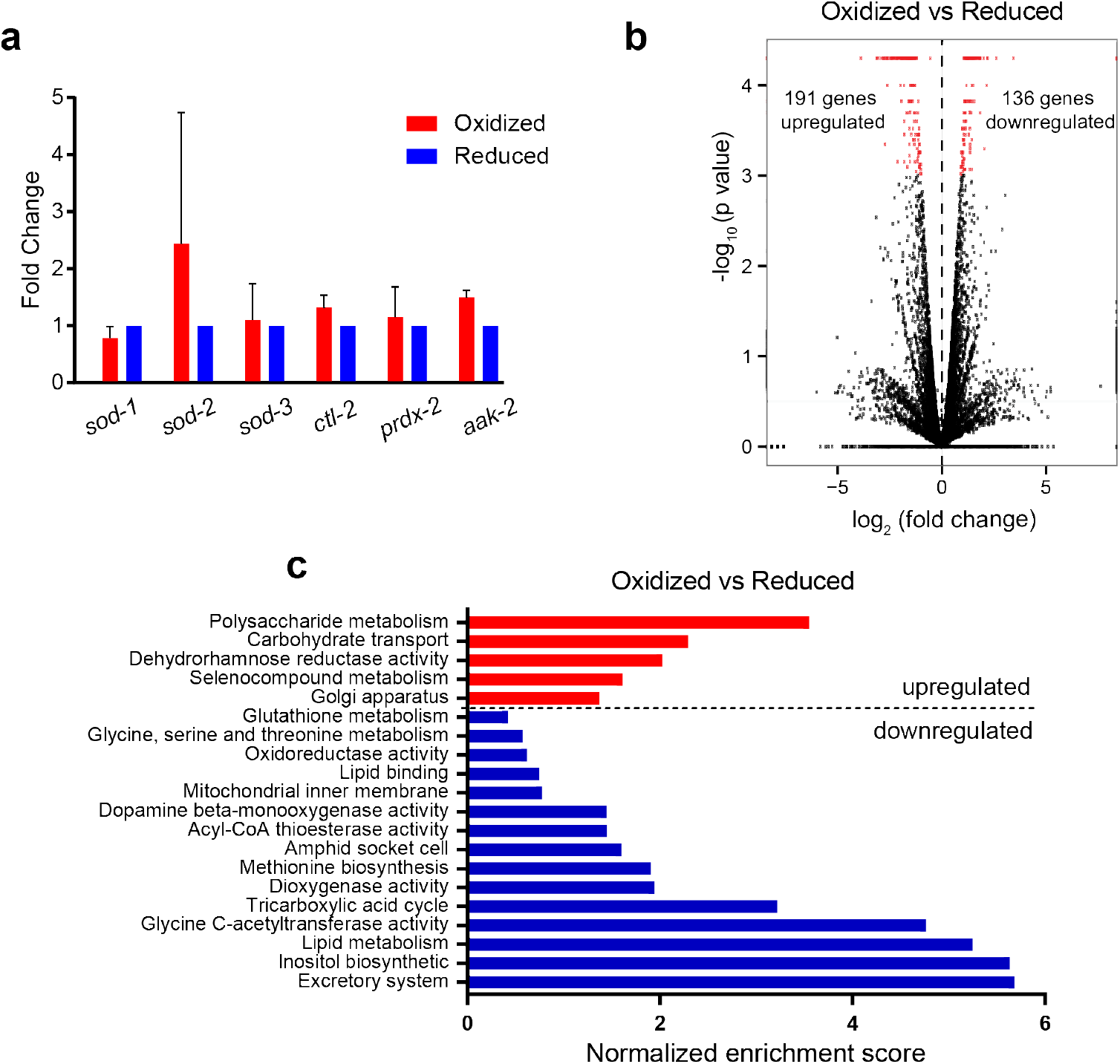
Gene expression profiles of oxidized and reduced subpopulations. (a) Steady-state transcript levels of selected oxidative stress-related genes of oxidized and reduced subpopulations at the L2 stage, as assessed by qRT-PCR, using *pmp-3* as reference gene. N≥3 experiments. No significant difference (paired *t*-test). Data were normalized to the reduced subpopulation and represent mean ± SEM. (b) Volcano plot showing fold changes versus p values for the transcriptomes of oxidized versus reduced subpopulations. Differentially expressed genes (DEGs, p≤0.05, changes ≥ log_2_ ± 0.6) are represented by red dots. Data were collected from 4 independent biological replicates of worms sorted at the L2 stage. (c) Gene Set Enrichment Analysis of the 327 DEGs in oxidized subpopulations compared to the reduced. Normalized enrichment scores (see methods for calculation) are represented by the bar graph. Terms (for summary, see Supplementary Table 1) indicating origin/process/phenotype associated with genes known to play a role in the process are shown on the left. Some terms (*) have been merged and are represented as a single category bar for simplicity (for detailed values, see Supplementary Table 1).

**Extended Data Figure 4.**
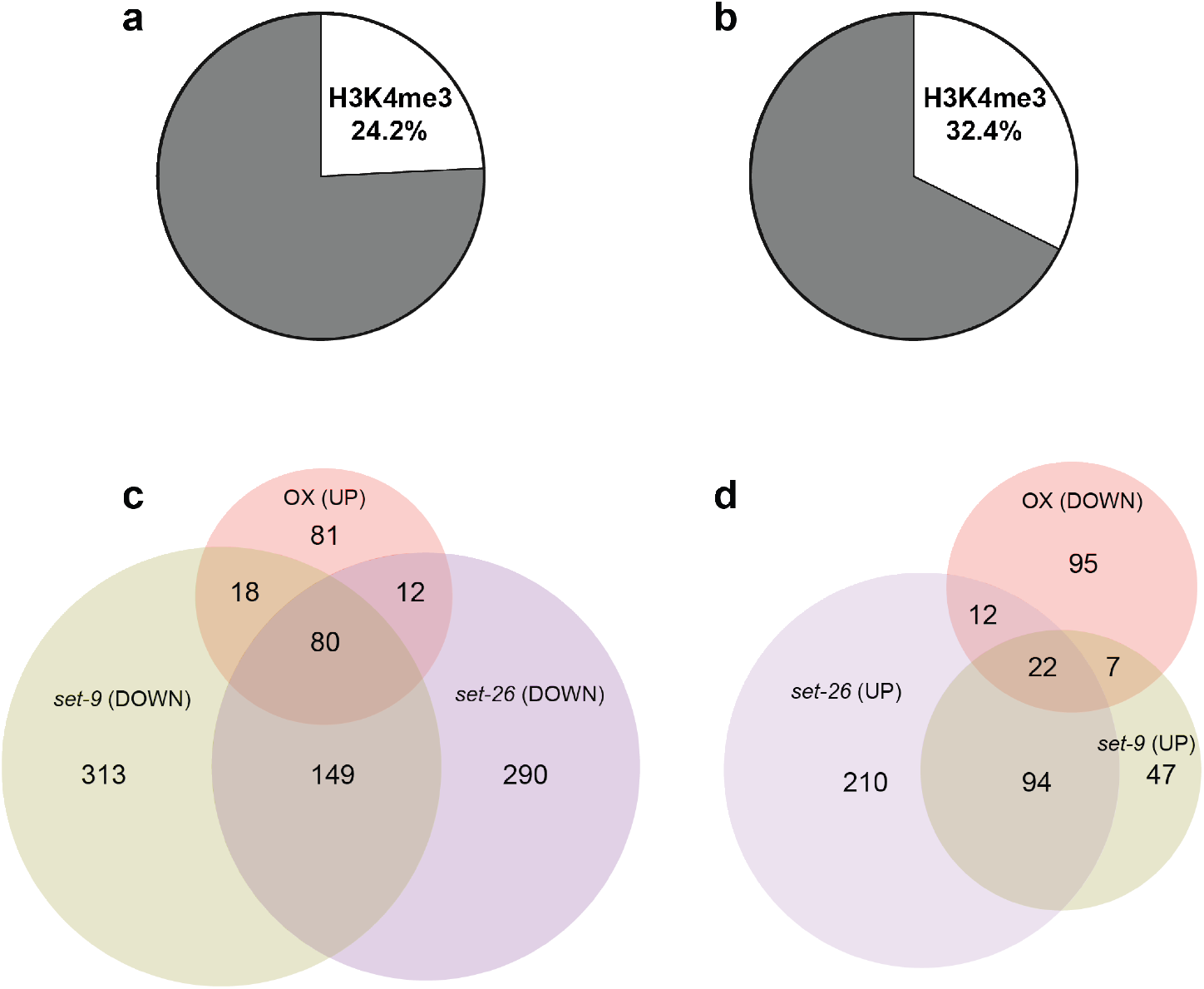
Comparison of DEGs from RNA-seq with publicly available H3K4me3 ChIP data sets. (a) and (b) The pie charts show the percentage of DEGs that intersect with H3K4me3 peak signals within their 5’ region (500 bp upstream and downstream from the transcription start site). The H3K4me3 ChIP data sets were generated from L3-staged N2 worms (ChIP–chip, GEO entry: GSE30789) in (a) and ChIP-seq, GEO entry: GSE28770) in (b). A significant overlap exists between differentially expressed genes in the oxidized subpopulation of worms and genes associated with H3K4me3 marks. Hypergeometric probability for (a): *P* = 0.064 and (b): *P* = 2.786 x 10^−6^. (c) and (d) Venn diagrams show the overlap among up-regulated (c) or down-regulated (d) gene sets in the oxidized L2-staged subpopulation and down – or up – regulated *set-9(rw5)* and *set-26(tm2467)* gene sets (GEO entry: GSE100623).

**Extended Data Figure 5.**
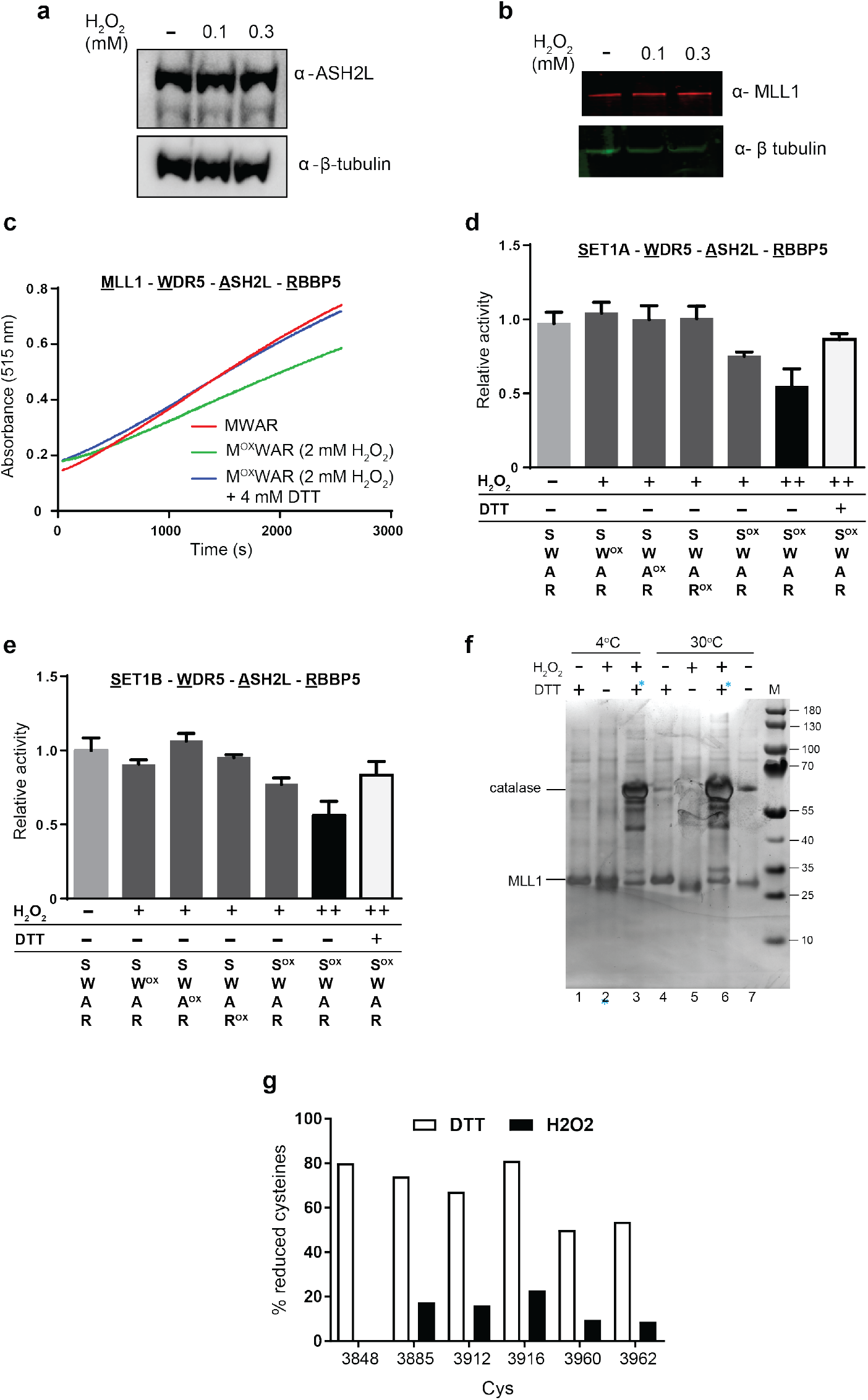
Histone methyltransferase complex activity and redox sensitivity. ASH2L (a) and MLL1 (b) levels after H2O2 treatment in HeLa cells, as assessed by western blot. For blot source images, see Supplementary Fig. 3. (c) Representative curves of the time course of the *in vitro* methyltransferase reaction for core HMC (MLL1-WDR5-ASH2L-RBBP5). Reaction rates were derived from the first 20 min of the linear range. (d, e) *In vitro* histone methyltransferase assay for core HMC containing GST-WR5, GST-ASH2L, GST-RBBP5 and either GST-SET1ASET (d) or GST-SET1BSET (e). Superscript OX indicates the protein that was pre-exposed to 1 mM (+) or 2 mM (++) H_2_O_2_ treatment before addition to the activity assay. DTT was added after the H_2_O_2_ treatment as indicated. N = 3 experiments for both (d) and (e). Data represent mean ± SEM. (f) MLL1_SET_ was treated with either 2 mM DTT, 2 mM H_2_O_2_ or 2 mM H_2_O_2_ followed by 4 mM DTT-treatment. Catalase was used to quench the H_2_O_2_. The proteins were denatured and thiols were modified with NEM prior to loading onto non-reducing SDS-PAGE to prevent non-specific thiol oxidation. The proteins were visualized by silver staining. M, marker. (g) Cysteine oxidation state in MLL1_SET_ after treatment with either 2 mM DTT or 2 mM H_2_O_2_ followed by NEM labeling as assessed by LC-MS/MS. The peptide containing Cys3967 could not be detected.

**Extended Data Figure 6.**
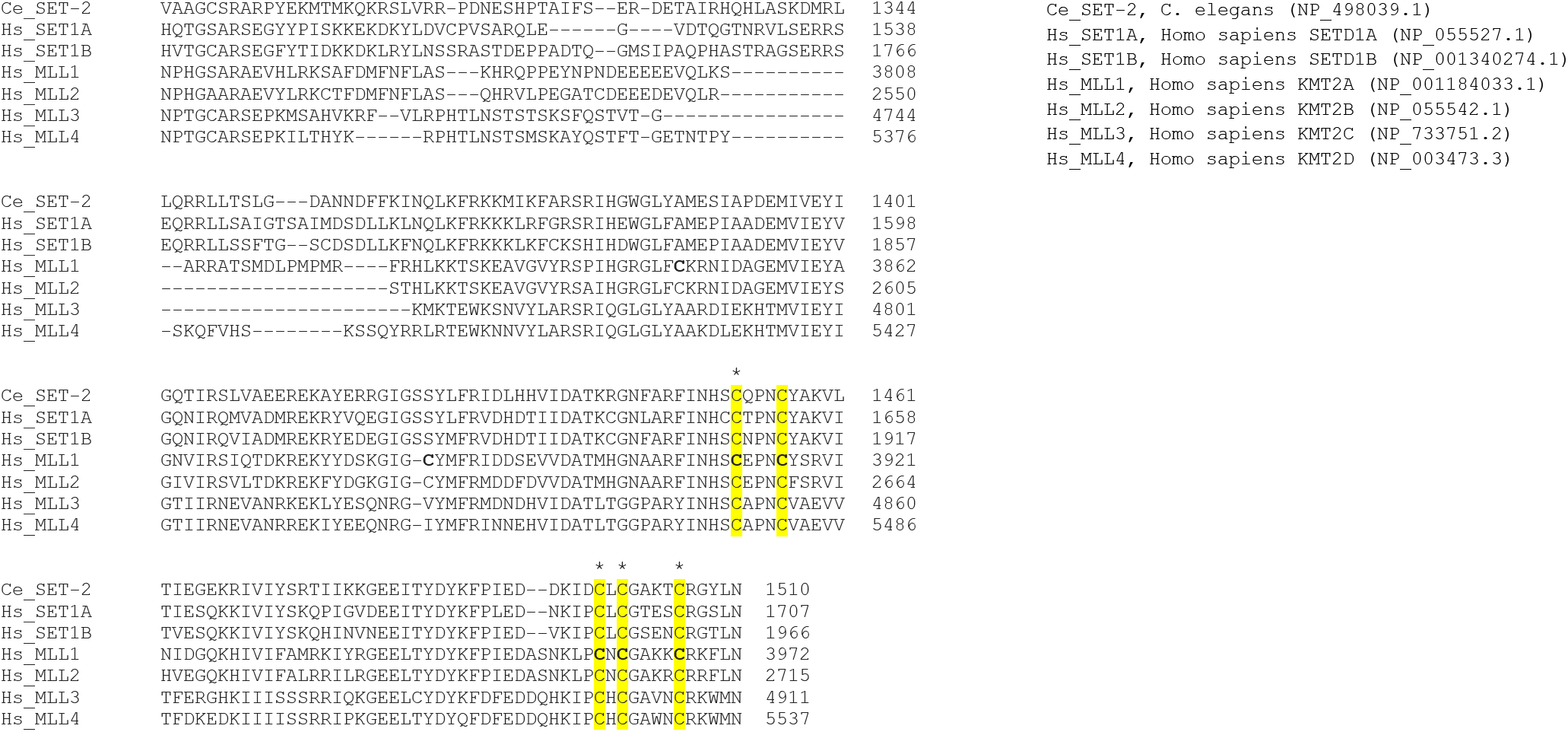
Sequence alignment of the SET-domain. All cysteines present in MLL1 are shown in bold, and the five absolutely conserved cysteines are highlighted in yellow. Cysteines shown to be involved in zinc coordination are marked with an asterisk. NCBI protein blast and Clustal Omega Multiple Sequence Alignment, Clustal O (1.2.4) were used.

**Extended Data Figure 7.**
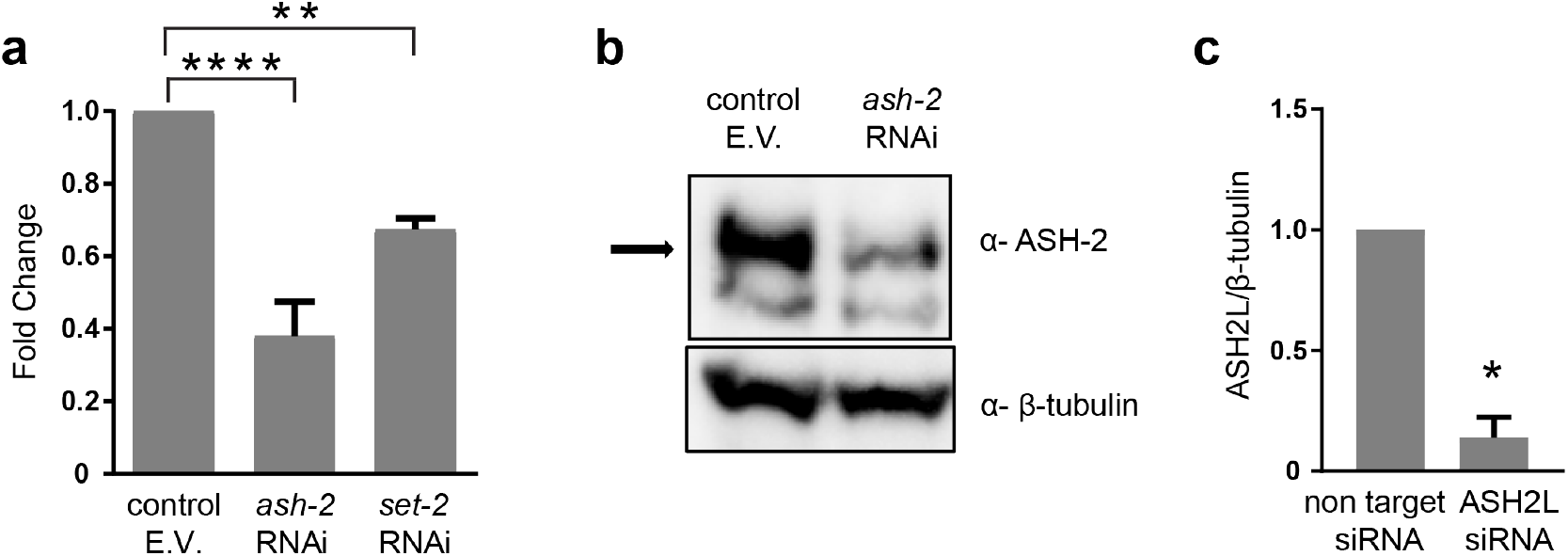
Knockdown efficiencies on components of H3K4me3 complex. (a) Transcript levels of N2*jrIs2*[P*rpl-17*::Grx1-roGFP2] worms treated with *ash-2* or *set-2* RNAi, as assessed by qRT-PCR, using *cdc-42* or panactin as reference genes. N = 6 (*ash-2*) and N = 2 (*set-2*) experiments. **, *p*<0.01; ****, *p*<0.0001 (unpaired *t*-test). E.V., empty vector. Data were normalized to the empty vector RNAi control. (b) Representative western blot analyzing ASH-2 levels in N2*jrls2*[P*rpl-17*::Grx1-roGFP2] worms treated with control RNAi or *ash-2* RNAi for 2 generations. For blot source image, see Supplementary Fig. 3. (c) ASH2L levels following *ash-2* siRNA treatment of HeLa cells. N = 2 experiments *, *p*< 0.05, (unpaired *t*-test). Data in (a) and (c) represent mean ± SEM.

**Extended Data Figure 8.**
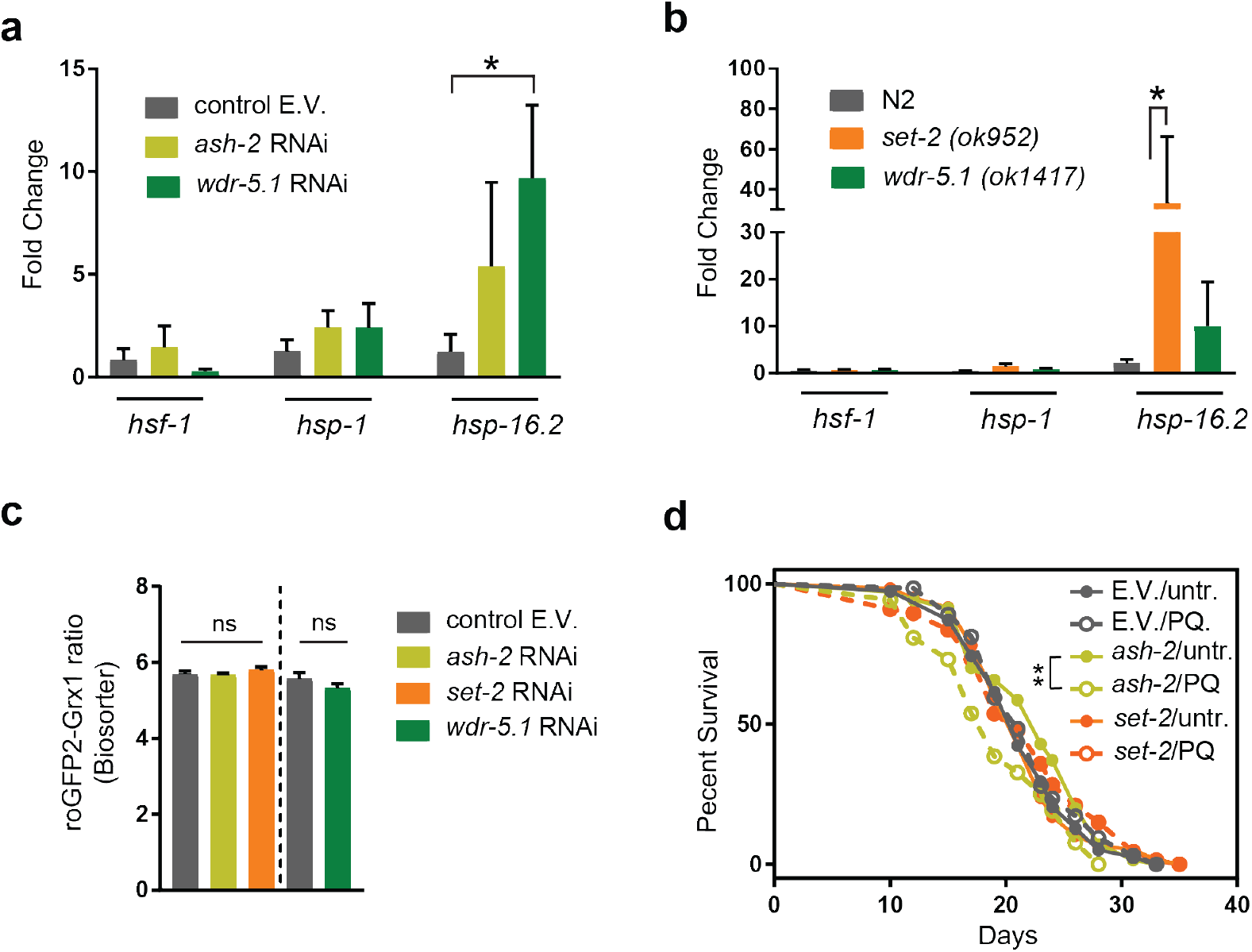
Effects of H3K4me3 down-regulation on heat shock response and endogenous redox state. (a) Transcript levels of selected heat shock genes after heat shock treatment of *N2jrIs2[Prpl-17::Grx1-roGFP2]* worms treated with the indicated RNAi. Data were normalized to the non-heat shock control and represent mean ± SEM. (b) Transcript levels of selected heat shock genes in *set-2* or *wdr-5.1* mutants before and after heat shock treatment. Heat shock and steady state data were normalized to the non-heat shock control and N2 worms, respectively and represent mean ± SEM. (a, b) N = 3 experiments. *, *p*<0.05 (one-way ANOVA with Bonferroni correction). Transcript levels were assessed by qRT-PCR, using *cdc-42* or spiked-in luciferase RNA as reference. *ns*, not significant (c) Grx1-roGFP2 ratios of L2 larval worms treated with *ash-2* RNAi, *set-2* RNAi, *wdr5.1* RNAi or the empty vector were measured using the BioSorter. N = 4 (*ash-2, set-2*) and N = 3 (*wdr-5.1*) experiments. All data represent mean ± SEM. ns, not significant (unpaired *t*-test) (d) Representative survival curves of *N2jrls2[Prpl-17*::Grx1-roGFP2] worms treated with *ash-2* or *set-2* RNAi for 2 generations, and treated with 1 mM PQ for 10 hours at the L2 larval stage. **, *p*<0.01 (log rank test). For lifespan values and n numbers, see Extended Data Table 4.

**Extended Data Table 1.** Lifespan assays of oxidized, mean and reduced subpopulations following heat shock treatment or in the continuous presence of paraquat or juglone.

| Condition | Experiment | Redox status | Mean | Max | Dead/ censored worms | *p* value | % change | Figure on text | Ratio BioSorter<sup>‡</sup> | *p* value (BioS.) |
| --- | --- | --- | --- | --- | --- | --- | --- | --- | --- | --- |
| Heat shock (long term) | 1 | Oxidized | 19 | 32.5 | 24 |  |  |  | 7.44±0.01(n=238) |  |
|  |  | Mean | 17 | 31.5 | 19/4 | 0.22 | 10.5%* | Figure 2b | 7.37±0.02(n=135) | 0.002 |
|  |  | Reduced | 13 | 27.3 | 32/1 | 0.0088 | 31.6%* |  | 7.29±0.02(n=132) | <0.0001 |
|  | 2 | Oxidized | 17.5 | 25 | 29/13 |  |  |  | 7.06±0.01(n=698) |  |
|  |  | Reduced | 12 | 21 | 20/3 | 0.0077 | 31.4%* |  | 7.01±0.012(n=319) | 0.002 |
|  | 3 | Oxidized | 18 | 25.9 | 96/2 |  |  |  | 7.15±0.007(n=893) |  |
|  |  | Reduced | 15 | 24.2 | 77/1 | 0.0153 | 16.7%* |  | 7.12±0.012(n=914) | 0.009 |
| Paraquat (2 mM) | 1 | Oxidized | 17 | 26 | 13 |  |  |  | 7.09±0.02(n=116) |  |
|  |  | Mean | 14 | 23.5 | 11 | 0.5818 | 17.6%* | Figure 2c | 6.99±0.033(n=113) | 0.006 |
|  |  | Reduced | 10 | 21 | 20 | 0.0215 | 41.2%* |  | 6.67±0.045(n=108) | <0.0001 |
|  | 2 | Oxidized | 13 | 18.6 | 38 |  |  |  | 7.13±0.03(n=32) |  |
|  |  | Reduced | 11 | 18.5 | 40 | 0.0456 | 15.4%* |  | 7.13±0.03(n=32) | <0.0001 |
|  | 3 | Oxidized | 13 | 19 | 47 |  |  |  | 7.35±0.017(n=158) |  |
|  |  | Reduced | 12 | 15.6 | 38 | 0.0037 | 17.9%<sup>†</sup> |  | 7.14±0.011(n=277) | <0.0001 |
| Juglone (250 uM) | 1 | Oxidized | 23 | 32.4 | 72 |  |  |  | 6.59±0.012(n=428) |  |
|  |  | Reduced | 19 | 32.4 | 73 | 0.0033 | 17.4%* |  | 6.55±0.012(n=513) | 0.007 |
|  | 2 | Oxidized | 18 | 33.3 | 86 |  |  |  | 7.27±0.01(n=161) |  |
|  |  | Reduced | 18 | 29.5 | 59 | 0.0202 | 11.4%<sup>†</sup> |  | 7.14±0.011(n=277) | <0.0001 |

Worms were sorted into the indicated subpopulations at the L2 stage, collected into NGM plates and exposed to heat shock treatment or to the indicated compounds. *p* values for Kaplan Meier survival analysis were calculated based on the log-rank (Mantel-Cox) method. Samples were compared to oxidized subpopulations. Maximum lifespan was defined as the 10% of last survival population.
\*, percentage change based on mean lifespan.
<sup>†</sup>, percentage change based on max lifespan.
<sup>‡</sup>, Grx1-roGFP2 ratio of subpopulations at the L2 stage, based on BioSorter analysis during sorting.

**Extended Data Table 2.** Lifespan assays of oxidized, mean and reduced subpopulations.

| Exp. | Redox status | Mean | Max | Dead/<br>censored<br>worms | *p*<br>value<br>(survival) | %<br>change | Figure<br>on<br>text | Ratio<br>BioSorter<sup>§</sup> | *p*<br>value<br>(BioS.) | Ratio<br>Microscope | *p* value<br>(Micr.) |
| --- | --- | --- | --- | --- | --- | --- | --- | --- | --- | --- | --- |
| 1 | Oxidized | 30 | 35.8 | 97/5 |  |  |  | 6.88±0.02(n=133) |  | 0.49±0.008(n=36) |  |
|  | Mean | 26 | 33.7 | 74/2 | 0.007 | 13.3%* | Fig. 2d | 6.69±0.02(n=143) | <0.0001 | 0.45±0.003(n=49) | <0.0001 |
|  | Reduced | 25 | 32.3 | 58/3 | 0.0004 | 17%* |  | 6.62±0.04(n=131) | <0.0001 | 0.44±0.003(n=50) | <0.0001 |
| 2 | Oxidized | 28 | 35.1 | 78/2 |  |  |  | 7.14±0.02(n=127) |  | 0.96±0.07(n=7) |  |
|  | Mean | 25 | 34.3 | 105/3 | 0.1018 | 10.7%* |  | 7.11±0.03(n=133) | 0.532 | 0.69±0.01(n=7) | 0.0027 |
|  | Reduced | 23 | 33.5 | 63/1 | 0.0118 | 18%* |  | 6.94±0.07(n=136) | 0.0101 | 0.66±0.01(n=10) | 0.002 |
| 3 | Oxidized | 21 | 34 | 94/2 |  |  |  | 7.45±0.02(n=139) |  | 0.51±0.03(n=14) |  |
|  | Reduced | 20 | 30 | 51 | 0.0088 | 12%<sup>†</sup> |  | 7.35±0.01(n=716) | 0.0003 | 0.4±0.01(n=13) | 0.0036 |
| 4 | Oxidized | 31 | 39.4 | 162/3 |  |  |  | 6.61±0.02(n=170) |  | 0.6±0.02(n=48) |  |
|  | Reduced | 28.5 | 38.8 | 120 | 0.042 | 8%* |  | 6.55±0.02(n=161) | 0.0456 | 0.54±0.01(n=44) | 0.0037 |
| 5 | Oxidized | 28 | 38 | 114/4 |  |  |  | 7.29±0.01(n=191) |  | 0.48±0.03(n=10) |  |
|  | Reduced | 26 | 35.4 | 106 | 0.0356 | 7%* |  | 7.12±0.02(n=88) | <0.0001 | 0.39±0.003(n=14) | 0.063 |
| 6<sup>¶</sup> | Oxidized | 24 | 34 | 164/7 |  |  |  | 6.19±0.02(n=275) |  | 1.68±0.01(n=9) |  |
|  | Reduced | 23 | 34 | 155/9 | 0.0862<sup>‡</sup> | 4%* |  | 6.03±0.02(n=273) | <0.0001 | 1.646±0.01(n=8) | 0.058 |
| 7 | E.V.oxidized | 24 | 36 | 48 |  |  | Figure 5d | 6.37±0.035(n=60) |  |  |  |
|  | E.V.reduced | 21 | 28.3 | 63 | 0.0004 | 12.5%* |  | 5.85±0.076(n=63) | <0.0001 |  |  |
| 8 | E.V.oxidized | 24 | 33 | 77/3 |  |  | Figure 5e | 5.55±0.04(n=114) |  |  |  |
|  | E.V.reduced | 20 | 32.7 | 80 | 0.002 | 17%* |  | 4.49±0.067(n=114) | <0.0001 |  |  |
| 9 | E.V.oxidized | 20 | 33.25 | 85 |  |  |  | 6.08±0.046(n=133) |  |  |  |
|  | E.V.reduced | 17 | 30.7 | 77 | 0.0264 | 15%* |  | 5.87±0.052(n=107) | 0.0032 |  |  |
| 10 | E.V.oxidized | 18 | 28.7 | 89/1 |  |  |  | 6.03±0.022(n=225) |  |  |  |
|  | E.V.reduced | 18 | 25.4 | 85 | 0.0077 | 11.5%<sup>†</sup> |  | 5.10±0.038(n=226) | <0.0001 |  |  |

\*, percentage change based on mean lifespan.
<sup>†</sup>, percentage change based on max lifespan.
<sup>‡</sup>, *p* value for survival analysis calculated based on the Gehan-Breslow-Wilcoxon method.
<sup>§</sup>, Grx1-roGFP2 ratio of subpopulations at the L2 stage, based on BioSorter analysis during sorting.
<sup>||</sup>, Grx1-roGFP2 ratio of subpopulations based on microscopic analysis at the L2 stage, following sorting. <sup>¶</sup>, Lifespan conducted with experimenter blind to the type of worms assayed.

**Extended Data Table 3.**
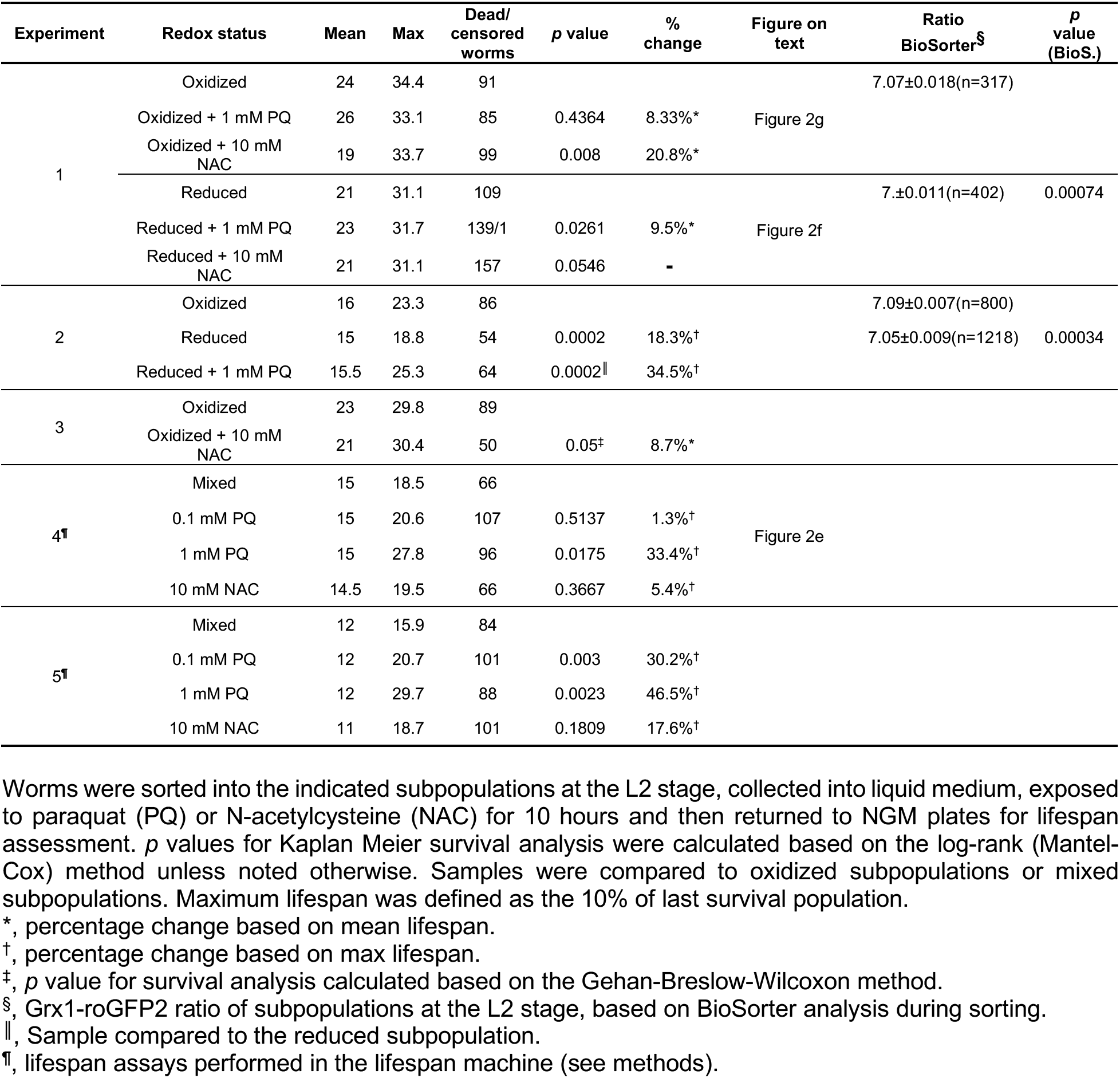
Lifespan assays of L2 subpopulations following transient exposure to oxidizing or reducing conditions.

**Extended Data Table 4.**
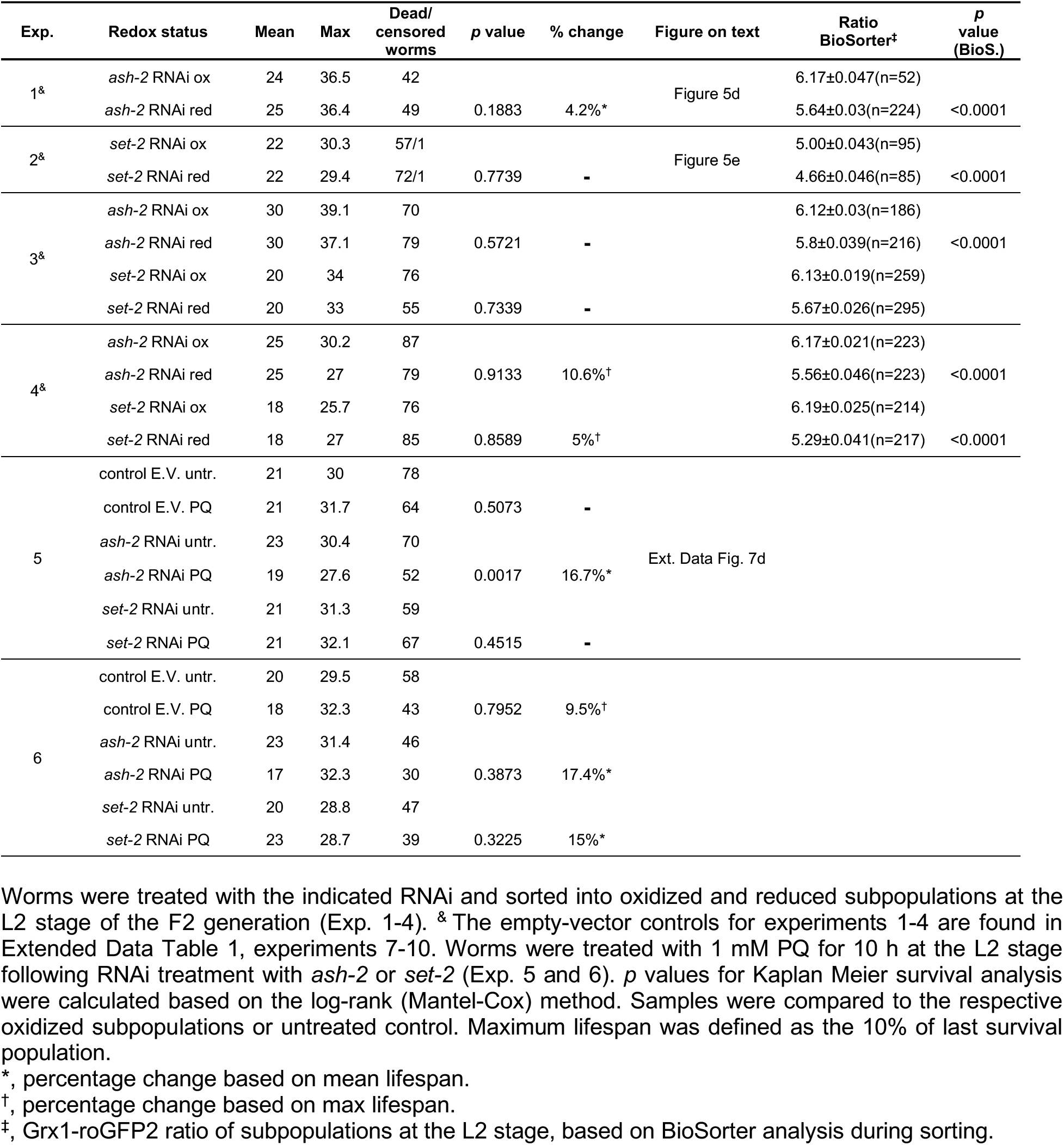
Lifespan assays upon H3K4me3-targeting RNAi treatment.

| Exp. | Redox status | Mean | Max | Dead/<br>censored<br>worms | p value | % change | Figure on text | Ratio<br>BioSorter<sup>‡</sup> | p<br>value<br>(BioS.) |
| --- | --- | --- | --- | --- | --- | --- | --- | --- | --- |
| 1<sup>&</sup> | *ash-2* RNAi ox | 24 | 36.5 | 42 |  |  | Figure 5d | 6.17±0.047(n=52) |  |
|  | *ash-2* RNAi red | 25 | 36.4 | 49 | 0.1883 | 4.2%* |  | 5.64±0.03(n=224) | <0.0001 |
| 2<sup>&</sup> | *set-2* RNAi ox | 22 | 30.3 | 57/1 |  |  | Figure 5e | 5.00±0.043(n=95) |  |
|  | *set-2* RNAi red | 22 | 29.4 | 72/1 | 0.7739 | - |  | 4.66±0.046(n=85) | <0.0001 |
| 3<sup>&</sup> | *ash-2* RNAi ox | 30 | 39.1 | 70 |  |  |  | 6.12±0.03(n=186) |  |
|  | *ash-2* RNAi red | 30 | 37.1 | 79 | 0.5721 | - |  | 5.8±0.039(n=216) | <0.0001 |
|  | *set-2* RNAi ox | 20 | 34 | 76 |  |  |  | 6.13±0.019(n=259) |  |
|  | *set-2* RNAi red | 20 | 33 | 55 | 0.7339 | - |  | 5.67±0.026(n=295) |  |
| 4<sup>&</sup> | *ash-2* RNAi ox | 25 | 30.2 | 87 |  |  |  | 6.17±0.021(n=223) |  |
|  | *ash-2* RNAi red | 25 | 27 | 79 | 0.9133 | 10.6%<sup>†</sup> |  | 5.56±0.046(n=223) | <0.0001 |
|  | *set-2* RNAi ox | 18 | 25.7 | 76 |  |  |  | 6.19±0.025(n=214) |  |
|  | *set-2* RNAi red | 18 | 27 | 85 | 0.8589 | 5%<sup>†</sup> |  | 5.29±0.041(n=217) | <0.0001 |
| 5 | control E.V. untr. | 21 | 30 | 78 |  |  | Ext. Data Fig. 7d |  |  |
|  | control E.V. PQ | 21 | 31.7 | 64 | 0.5073 | - |  |  |  |
|  | *ash-2* RNAi untr. | 23 | 30.4 | 70 |  |  |  |  |  |
|  | *ash-2* RNAi PQ | 19 | 27.6 | 52 | 0.0017 | 16.7%* |  |  |  |
|  | *set-2* RNAi untr. | 21 | 31.3 | 59 |  |  |  |  |  |
|  | *set-2* RNAi PQ | 21 | 32.1 | 67 | 0.4515 | - |  |  |  |
| 6 | control E.V. untr. | 20 | 29.5 | 58 |  |  |  |  |  |
|  | control E.V. PQ | 18 | 32.3 | 43 | 0.7952 | 9.5%<sup>†</sup> |  |  |  |
|  | *ash-2* RNAi untr. | 23 | 31.4 | 46 |  |  |  |  |  |
|  | *ash-2* RNAi PQ | 17 | 32.3 | 30 | 0.3873 | 17.4%* |  |  |  |
|  | *set-2* RNAi untr. | 20 | 28.8 | 47 |  |  |  |  |  |
|  | *set-2* RNAi PQ | 23 | 28.7 | 39 | 0.3225 | 15%* |  |  |  |

\*, percentage change based on mean lifespan.
<sup>†</sup>, percentage change based on max lifespan.
<sup>‡</sup>, Grx1-roGFP2 ratio of subpopulations at the L2 stage, based on BioSorter analysis during sorting.

## ACKNOWLEDGEMENTS

We are highly indebted to M. Malinouski and T. Mullins from Union Biometrica for their invaluable help and assistance with the reconfiguration of the Biosorter. We thank G. Csankovszki for antibodies, *C. elegans* RNAi feeding clones and comments, B. Braeckman for the N2*jrIs2*[P*rpl-17::*Grx-1-roGFP2] strain, J. Nandakumar for HeLa (EM-2-11ht) cells, the *Caenorhabditis* Genetics Center (funded by National Institutes of Health Infrastructure Program P40 OD010440) for strains, the DNA Sequencing Core (BRCF), R. Tagett and the Bioinformatics Core of University of Michigan for RNA sequencing, M. Kim for setting up the lifespan machine, K. Wan for protein purification and Jakob laboratory members for comments on the manuscript. We also thank R. Sawarkar and J. Labbadia for important suggestions, and J. Bardwell for critically reading the manuscript. Mass spectrometry was performed by MS Bioworks. This work was supported by NIH grants GM122506 and AG046799 as well as the Priority Program SPP 1710 of the Deutsche Forschungsgemeinschaft (Schw823/3-2) to U.J., a NIH T32 Career Training in the Biology of Aging grant to D.B. and B.O., and the National Natural Science Foundation of China (31470737) to Y.C.

## AUTHOR CONTRIBUTIONS

**D.B.** Conceived and conducted most experiments, data analysis, and wrote ms.

**D.K.** Conceived and conducted experiments

**Y.Z.** Conducted experiments

**K.U.** Conducted experiments

**B.O.** Conducted experiments

**L.X.** Conducted experiments

**A.K.** Conducted experiments

**Y-T. L.** Conducted experiments

**Y.D.** Conceived experiments and provided material

**S.Q.** Conceived experiments

**Y.C.** Conceived experiments

**U.J.** Conceived experiments, conducted data analysis, and wrote the ms.

## AUTHOR INFORMATION

Reprints and permissions information is available at www.nature.com/reprints

The authors declare no competing financial interests.

Correspondence and requests for materials should be addressed to

## References

1 Bladbjerg, E. M. et al. Genetic influence on thrombotic risk markers in the elderly--a Danish twin study. Journal of thrombosis and haemostasis: JTH 4, 599–607, doi:10.1111/j.1538-7836.2005.01778.x (2006).

2 Herskind, A. M. et al. The heritability of human longevity: a population-based study of 2872 Danish twin pairs born 1870-1900. Human genetics 97, 319–323 (1996).

3 Finch, C. E. & Kirkwood, T. B. Chance, Development and Aging. (Oxford University Press, 2000).

4 Finch, C. E. & Tanzi, R. E. Genetics of aging. Science (New York, N.Y 278, 407–411 (1997).

5 Kirkwood, T. B. & Finch, C. E. Ageing: the old worm turns more slowly. Nature 419, 794–795 (2002).

6 Rea, S. L., Wu, D., Cypser, J. R., Vaupel, J. W. & Johnson, T. E. A stress-sensitive reporter predicts longevity in isogenic populations of Caenorhabditis elegans. Nature genetics 37, 894–898 (2005).

7 Cross, J. V. & Templeton, D. J. Regulation of signal transduction through protein cysteine oxidation. Antioxidants & redox signaling 8, 1819–1827 (2006).

8 D’Autreaux, B. & Toledano, M. B. ROS as signalling molecules: mechanisms that generate specificity in ROS homeostasis. Nature reviews 8, 813–824 (2007).

9 Holmstrom, K. M. & Finkel, T. Cellular mechanisms and physiological consequences of redox-dependent signalling. Nature reviews 15, 411–421, doi:10.1038/nrm3801 (2014).

10 Lee, S. J., Hwang, A. B. & Kenyon, C. Inhibition of respiration extends C. elegans life span via reactive oxygen species that increase HIF-1 activity. Current biology: CB 20, 2131–2136, doi:10.1016/j.cub.2010.10.057 (2010).

11 Schulz, T. J. et al. Glucose restriction extends Caenorhabditis elegans life span by inducing mitochondrial respiration and increasing oxidative stress. Cell metabolism 6, 280–293, doi:10.1016/j.cmet.2007.08.011 (2007).

12 Dillin, A. et al. Rates of behavior and aging specified by mitochondrial function during development. Science (New York, N.Y 298, 2398–2401 (2002).

13 Desjardins, D. et al. Antioxidants reveal an inverted U-shaped dose-response relationship between reactive oxygen species levels and the rate of aging in Caenorhabditis elegans. Aging cell 16, 104–112.

14 Heidler, T., Hartwig, K., Daniel, H. & Wenzel, U. Caenorhabditis elegans lifespan extension caused by treatment with an orally active ROS-generator is dependent on DAF-16 and SIR-2.1. Biogerontology 11, 183–195.

15 Yang, W. & Hekimi, S. A mitochondrial superoxide signal triggers increased longevity in Caenorhabditis elegans. PLoS biology 8, e1000556.

16 Ristow, M. & Schmeisser, S. Extending life span by increasing oxidative stress. Free radical biology & medicine 51, 327–336, doi:10.1016/j.freeradbiomed.2011.05.010 (2011).

17 Knoefler, D. et al. Quantitative in vivo redox sensors uncover oxidative stress as an early event in life. Molecular cell 47, 767–776.

18 Gutscher, M. et al. Real-time imaging of the intracellular glutathione redox potential. Nature methods 5, 553–559 (2008).

19 Back, P. et al. Exploring real-time in vivo redox biology of developing and aging Caenorhabditis elegans. Free radical biology & medicine 52, 850–859.

20 Labbadia, J. & Morimoto, R. I. Repression of the Heat Shock Response Is a Programmed Event at the Onset of Reproduction. Molecular cell 59, 639–650.

21 Morton, E. A. & Lamitina, T. Caenorhabditis elegans HSF-1 is an essential nuclear protein that forms stress granule-like structures following heat shock. Aging cell 12, 112–120.

22 Greer, E. L. et al. Members of the H3K4 trimethylation complex regulate lifespan in a germline-dependent manner in C. elegans. Nature 466, 383–387.

23 Dou, Y. et al. Regulation of MLL1 H3K4 methyltransferase activity by its core components. Nature structural & molecular biology 13, 713–719, doi:10.1038/nsmb1128 (2006).

24 Shilatifard, A. The COMPASS family of histone H3K4 methylases: mechanisms of regulation in development and disease pathogenesis. Annual review of biochemistry 81, 65–95, doi:10.1146/annurev-biochem-051710-134100 (2012).

25 Guenther, M. G., Levine, S. S., Boyer, L. A., Jaenisch, R. & Young, R. A. A chromatin landmark and transcription initiation at most promoters in human cells. Cell 130, 77–88, doi:10.1016/j.cell.2007.05.042 (2007).

26 Wang, W. et al. SET-9 and SET-26 are H3K4me3 readers and play critical roles in germline development and longevity. eLife 7, doi:10.7554/eLife.34970 (2018).

27 Cao, F. et al. Targeting MLL1 H3K4 methyltransferase activity in mixed-lineage leukemia. Molecular cell 53, 247–261, doi:10.1016/j.molcel.2013.12.001 (2014).

28 Wu, M. et al. Molecular regulation of H3K4 trimethylation by Wdr82, a component of human Set1/COMPASS. Molecular and cellular biology 28, 7337–7344, doi:10.1128/MCB.00976-08 (2008).

29 Southall, S. M., Wong, P. S., Odho, Z., Roe, S. M. & Wilson, J. R. Structural basis for the requirement of additional factors for MLL1 SET domain activity and recognition of epigenetic marks. Molecular cell 33, 181–191 (2009).

30 Dharmarajan, V., Lee, J. H., Patel, A., Skalnik, D. G. & Cosgrove, M. S. Structural basis for WDR5 interaction (Win) motif recognition in human SET1 family histone methyltransferases. The Journal of biological chemistry 287, 27275–27289.

31 Patel, A., Dharmarajan, V., Vought, V. E. & Cosgrove, M. S. On the mechanism of multiple lysine methylation by the human mixed lineage leukemia protein-1 (MLL1) core complex. The Journal of biological chemistry 284, 24242–24256 (2009).

32 Patel, A., Vought, V. E., Dharmarajan, V. & Cosgrove, M. S. A conserved arginine-containing motif crucial for the assembly and enzymatic activity of the mixed lineage leukemia protein-1 core complex. The Journal of biological chemistry 283, 32162–32175 (2008).

33 Li, T. & Kelly, W. G. A role for Set1/MLL-related components in epigenetic regulation of the Caenorhabditis elegans germ line. PLoS Genet 7, e1001349, doi:10.1371/journal.pgen.1001349 (2011).

34 Allen, M. D. et al. Solution structure of the nonmethyl-CpG-binding CXXC domain of the leukaemia-associated MLL histone methyltransferase. The EMBO journal 25, 4503–4512, doi:10.1038/sj.emboj.7601340 (2006).

35 Cosgrove, M. S. & Patel, A. Mixed lineage leukemia: a structure-function perspective of the MLL1 protein. The FEBS journal 277, 1832–1842, doi:10.1111/j.1742-4658.2010.07609.x (2010).

36 Leichert, L. I. et al. Quantifying changes in the thiol redox proteome upon oxidative stress in vivo. Proceedings of the National Academy of Sciences of the United States of America 105, 8197–8202 (2008).

37 Weiner, A. et al. Systematic dissection of roles for chromatin regulators in a yeast stress response. PLoS biology 10, e1001369, doi:10.1371/journal.pbio.1001369 (2012).

38 Kenyon, C. J. The genetics of ageing. Nature 464, 504–512, doi:10.1038/nature08980 (2010).

39 Hansen, M., Flatt, T. & Aguilaniu, H. Reproduction, fat metabolism, and life span: what is the connection? Cell metabolism 17, 10–19, doi:10.1016/j.cmet.2012.12.003 (2013).

40 Han, S. et al. Mono-unsaturated fatty acids link H3K4me3 modifiers to C. elegans lifespan. Nature 544, 185–190, doi:10.1038/nature21686 (2017).

41 Sun, L., Sadighi Akha, A. A., Miller, R. A. & Harper, J. M. Life-span extension in mice by preweaning food restriction and by methionine restriction in middle age. The journals of gerontology. Series A, Biological sciences and medical sciences 64, 711–722, doi:10.1093/gerona/glp051 (2009).

42 Brenner, S. The genetics of Caenorhabditis elegans. Genetics 77, 71–94 (1974).

43 Stroustrup, N. et al. The Caenorhabditis elegans Lifespan Machine. Nature methods 10, 665–670, doi:10.1038/nmeth.2475 (2013).

44 Koopman, M. et al. A screening-based platform for the assessment of cellular respiration in Caenorhabditis elegans. Nature protocols 11, 1798–1816, doi:10.1038/nprot.2016.106 (2016).

45 Langmead, B., Trapnell, C., Pop, M. & Salzberg, S. L. Ultrafast and memory-efficient alignment of short DNA sequences to the human genome. Genome biology 10, R25 (2009).

46 Trapnell, C., Pachter, L. & Salzberg, S. L. TopHat: discovering splice junctions with RNA-Seq. Bioinformatics (Oxford, England) 25, 1105–1111 (2009).

47 Subramanian, A. et al. Gene set enrichment analysis: a knowledge-based approach for interpreting genome-wide expression profiles. Proceedings of the National Academy of Sciences of the United States of America 102, 15545–15550 (2005).

48 Edgar, R., Domrachev, M. & Lash, A. E. Gene Expression Omnibus: NCBI gene expression and hybridization array data repository. Nucleic acids research 30, 207–210 (2002).

49 Weidenfeld, I. et al. Inducible expression of coding and inhibitory RNAs from retargetable genomic loci. Nucleic acids research 37, e50 (2009).

50 Ng, S. B. et al. Exome sequencing identifies MLL2 mutations as a cause of Kabuki syndrome. Nature genetics 42, 790–793, doi:10.1038/ng.646 (2010).

51 Dorgan, K. M. et al. An enzyme-coupled continuous spectrophotometric assay for S-adenosylmethionine-dependent methyltransferases. Analytical biochemistry 350, 249–255 (2006).

52 Southall, S. M., Cronin, N. B. & Wilson, J. R. A novel route to product specificity in the Suv4-20 family of histone H4K20 methyltransferases. Nucleic acids research 42, 661–671.

