## Supplementary figures and images for "Developmental Reactive Oxygen Species Target Histone Methylation to Individualize Stress resistance and Lifespan"

### Supplementary data

##### Supplementary Figure 1

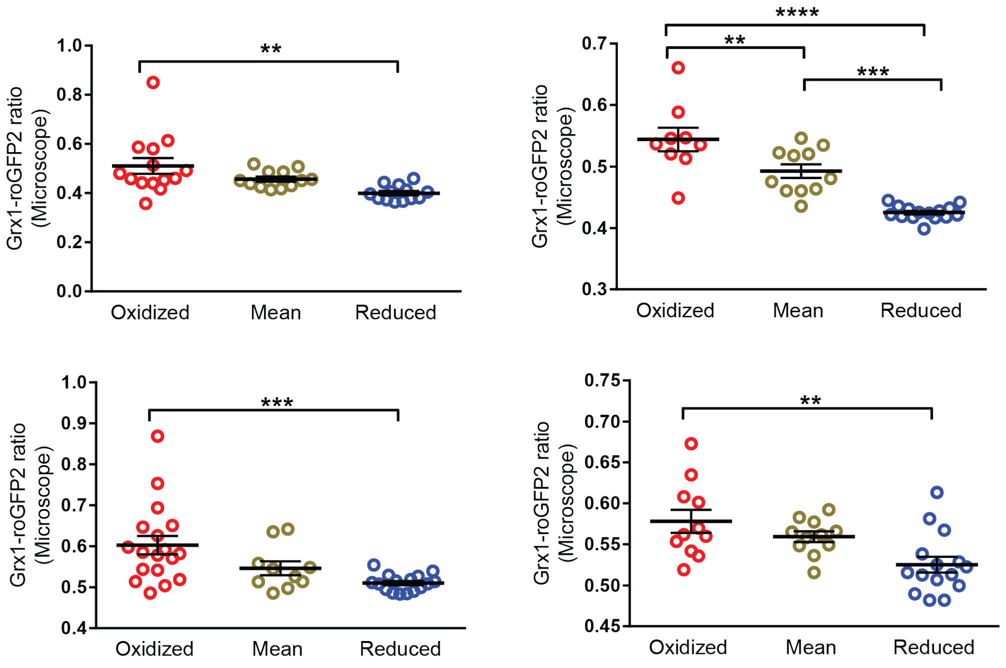

#### Supplementary Figure 2

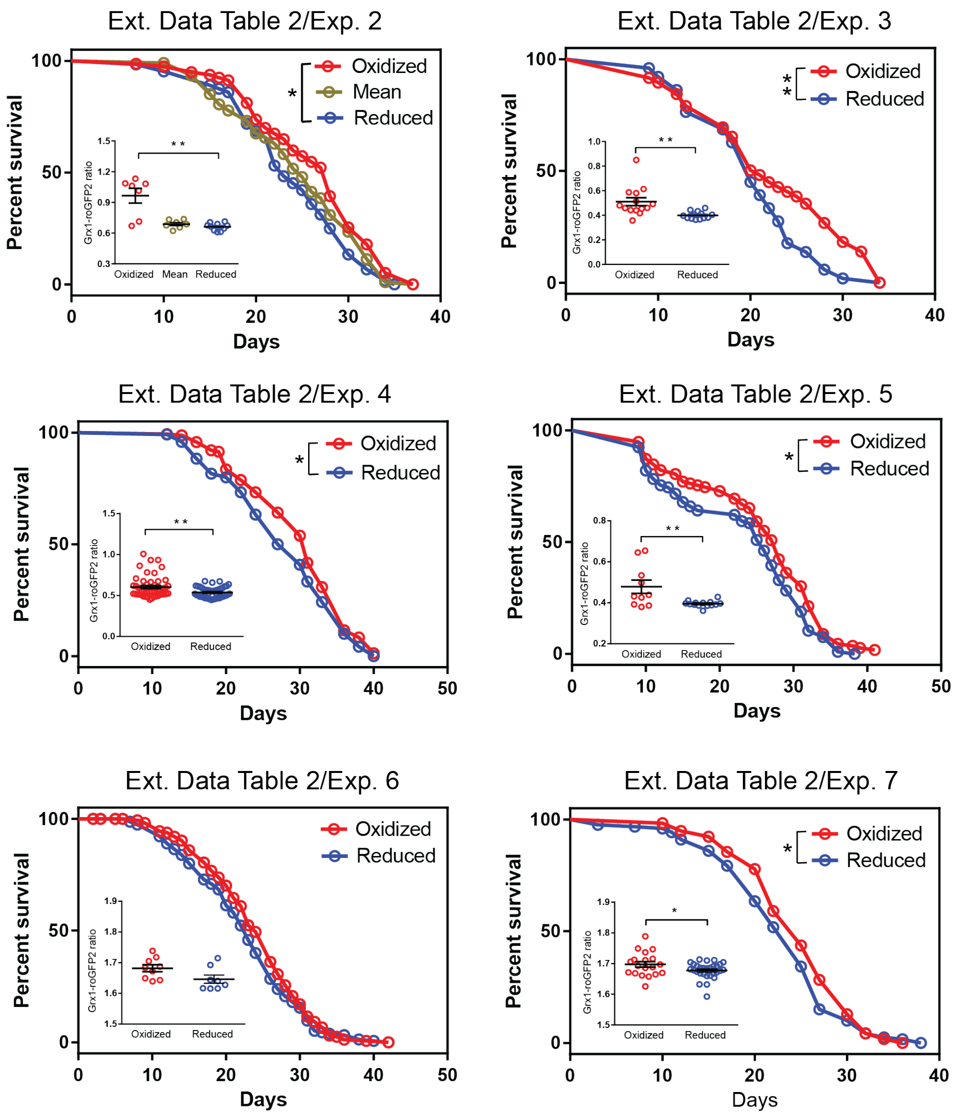

##### Supplementary Figure 3

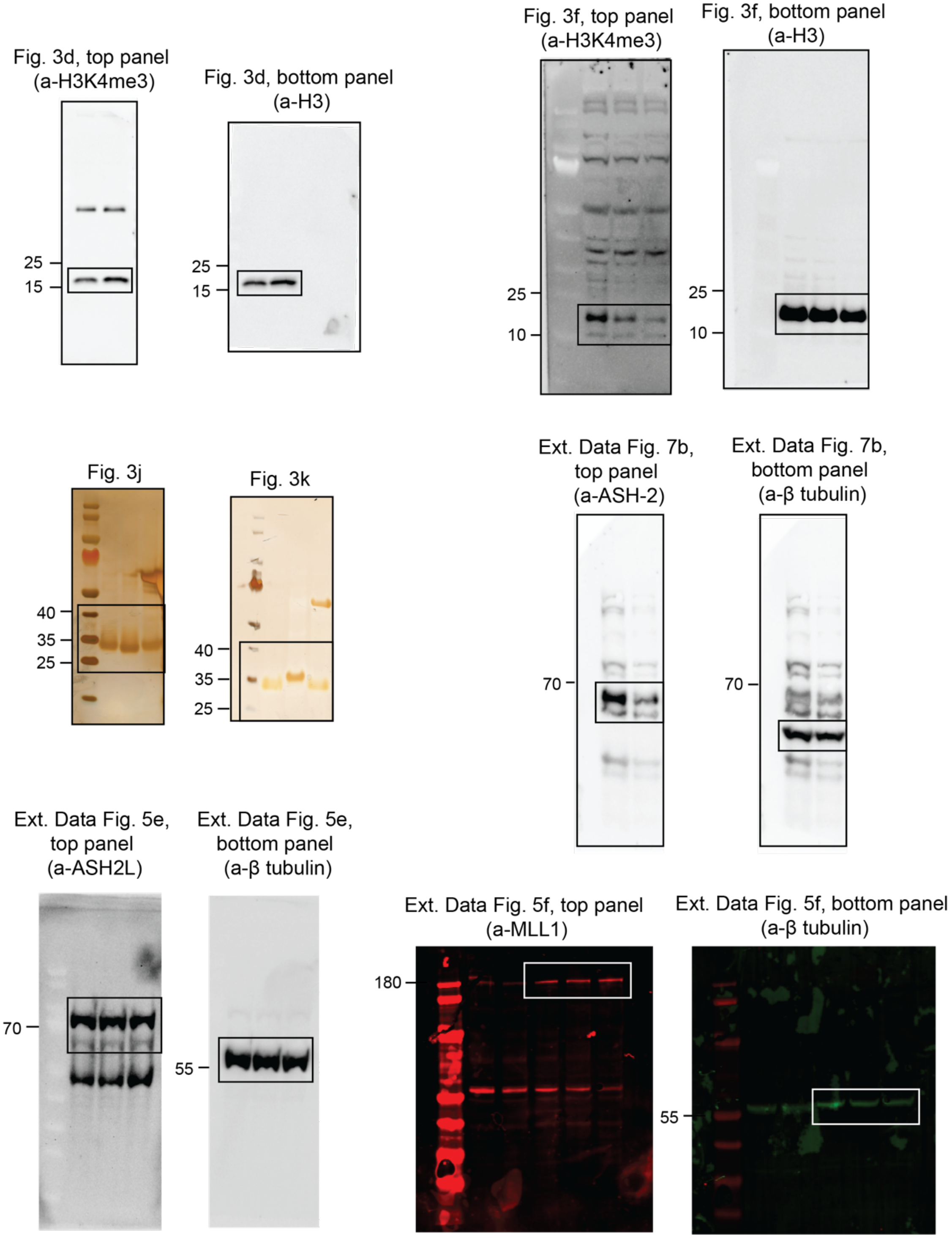
